# GBZ-base and GAF-base: Indexed pangenome file formats

**DOI:** 10.64898/2026.07.10.737775

**Authors:** Jouni Sirén, Benedict Paten, the Human Pangenome Reference Consortium

## Abstract

**Motivation:** Existing pangenome file formats are designed for batch processing. Graphs must be loaded into memory, and alignment files must be read sequentially. Indexed file formats that can be used directly from disk would be more appropriate for interactive applications.

**Results:** We propose GBZ-base and GAF-base — SQLite-backed file formats comparable to GBZ and GAF. GBZ-base supports efficient extraction of local subgraphs, and GAF-base lets us extract all alignments to the subgraph. Additionally, GAF-base is smaller than any other file format for sequence-to-graph alignments.

**Availability and implementation:** From https://github.com/jltsiren/gbz-base and https://crates.io/crates/gbz-base under the MIT license.

## Introduction

A *pangenome graph* represents an alignment of the underlying *haplotype sequences*. There is no single true pangenome graph for any given set of haplotypes. Instead, different alignments yield different graphs, which are useful for different purposes.

Common applications for pangenome graphs include read mapping (Garrison et al., 2018; Kim et al., 2019; Rautiainen and Marschall, 2020; Sirén et al., 2021, 2024; Chang et al., 2025) and variant calling (Kim et al., 2019; Hickey et al., 2020; Ebler et al., 2022; Asri et al., 2025). They improve accuracy over pipelines using a linear reference genome by reducing reference bias. This is particularly noticeable in regions, where the sequenced genome diverges substantially from the linear reference.

Method development for pangenomics is largely driven by human applications. Many tools come from the *Human Pangenome Reference Consortium* (HPRC) (Liao et al., 2023), which is building a human reference based on high-quality assemblies of hundreds of diverse genomes.

Existing pangenome file formats are designed for batch processing. Files containing graphs, indexes, or alignments must be typically either loaded into memory or read sequentially. This can be too slow or require too much memory in interactive tasks such as visualization, which are often run on end-user laptops. In this work, we propose indexed file formats, where we can quickly locate and read only the relevant parts of the pangenome.

### Pangenome graphs

Text-based *Graphical Fragment Assembly* (GFA) format is the primary interchange format for pangenome graphs. Because GFA represents each path explicitly, it scales poorly to graphs based on a large number of haplotypes. Grammar compression has been proposed to reduce the size of such GFA files while maintaining the simplicity and human-readability of the format (Heringer and Doerr, 2025).

GBZ (Sirén and Paten, 2022) is a pangenome graph file format used in the vg toolkit (Garrison et al., 2018) and related tools. It can handle graphs with a large number of haplotypes space-efficiently. Human GBZ graphs can be processed on a laptop, but loading the graph into memory still takes tens of seconds. While the format was designed with memory-mapping in mind, this turned out to be too impractical to implement.

We propose GBZ-base — an SQLite-backed file format that supports random access to the graph directly from disk. Since GBZ-base shares the data layout with GBZ, it removes the need for memory-mapped GBZ files. We can therefore develop new versions of the GBZ file format with better compression.

### Alignments

There are no established file formats for sequence-to-graph alignments. The vg toolkit uses its own GAM format internally. The *Graph Alignment Format* (GAF) introduced by minigraph (Li et al., 2020) is the closest to a de facto standard. However, the version of GAF written by the vg toolkit has diverged from the original specification (see Supplement 2 for details).

GAF is a text-based format that can be seen as the pangenome equivalent of SAM (Li et al., 2009). Each line contains an alignment record representing the alignment of an interval of the *query sequence* to an interval of the *target path*. The query sequence is not stored explicitly. Instead, there is a *difference string* describing the edit operations required to transform the target path to the query sequence. There is no binary version of GAF equivalent to BAM (Li et al., 2009) or a compressed version equivalent to CRAM (Fritz et al., 2011; Cochrane et al., 2013).

We propose GAF-base — an SQLite-backed binary format largely equivalent to GAF. As with indexed GAF files (Novak et al., 2024), we sort the alignments by intervals of node identifiers in the target path. We develop the approach further to extract alignments to any given subgraph. As in CRAM, we partition the alignments into blocks and use columnar compression within each block. The resulting file format is smaller than existing formats for sequence-to-graph alignments.

## Background

### Bidirected sequence graphs

Pangenome graphs are often represented as *bidirected sequence graphs G* = (*V, E, l*). We assume that *nodes v ∈ V* have integer *identifiers* (*V ⊂* N) and that identifier 0 is reserved for technical purposes. Each node *v ∈ V* has a non-empty sequence *label l*(*v*).

We can *visit* a node *v ∈ V* in the forward (left to right) and reverse (right to left) *orientations*, which we denote as 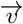 and 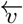, respectively. The label of the forward visit is 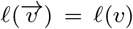, while the label of the reverse visit 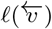 is its reverse complement. We give node visits integer identifiers called *handles* such that 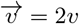 and 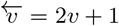 for all *v ∈ V*.

*Edges e ∈ E* are bidirectional and connect node visits. If *x* is a node visit, let 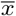 be the visit to the same node in the other orientation. If we have an edge (*x, y*) *∈ E*, we also have the reverse edge 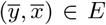. Bidirected graphs are often implemented as directed graphs *G*^*′*^ = (*V* ^*′*^, *E*^*′*^, *l*), where each node visit is a separate node. In other words, 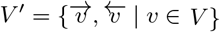 and *E*^*′*^ = *E*.

A *path P* = *x*_0_ *· · · x*_*m−*1_ is a sequence of node visits connected by edges. We have (*x*_*i*_, *x*_*i*+1_) *∈ E* for all 0 *≤ i < m −* 1. The label of path *P* is the concatenation *l*(*P*) = *l*(*x*_0_) *· · · l*(*x*_*m−*1_) of the labels of the visits. Reverse path *P* visits the same nodes in reverse order and the other orientation.

Its label 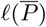 is therefore the reverse complement of *l*(*P*).

### GBZ file format

*GBZ* (Sirén and Paten, 2022) *is the pangenome graph file format used in the vg toolkit (Garrison et al*., *2018), when we want to store both the graph itself and the underlying haplotypes as paths. It is based on data structures used in the Giraffe aligner (Sirén et al*., *2021; Chang et al*., *2025)*. *Internally, a GBZ graph is a directed graph over the handles emulating a bidirected graph*.

*The key component of a GBZ graph is the GBWT index* (Sirén et al., 2020) storing the haplotype paths. Each path is encoded as a sequence of handles in both orientations and stored in an FM-index (Ferragina and Manzini, 2005). There are therefore two *stored paths* for each *original path*. The FM-index is partitioned between node records, one for each handle. Each record stores a list of outgoing edges and a list of path visits. See Figure 1 for an example. The records are compressed and concatenated, and an index is used for finding the record corresponding to a handle.

**Figure 1.**
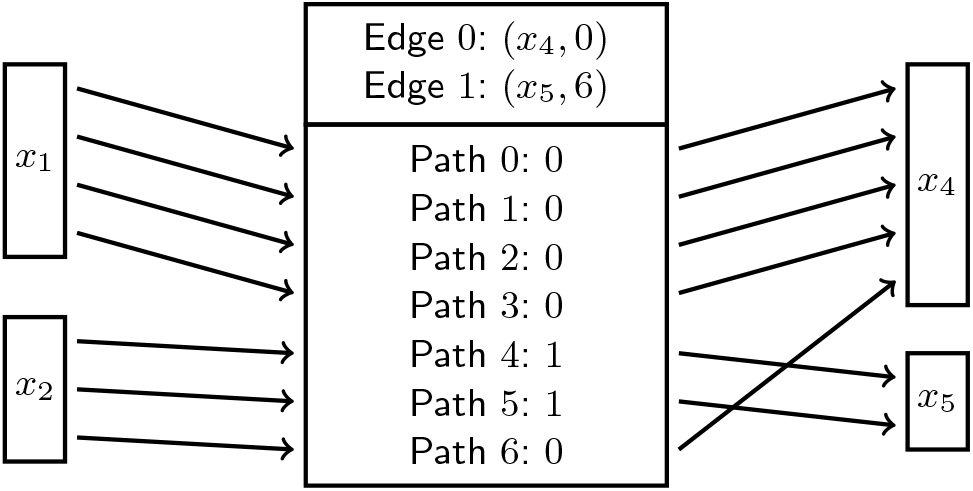
GBWT record for handle *x*_3_. Because *x*_1_ *< x*_2_, path visits coming from *x*_1_ are before those from *x*_2_ in colexicographic order. Each path visit is encoded as the rank of the outgoing edge. Nonzero offset in edge 1 indicates that there are path visits coming from handles *x < x*_3_ to handle *x*_5_.

A node record stores the path visits in colexicographic order by the prefix of the path up to the corresponding oriented node. For each path visit, we write the next handle in the path, encoding it as the rank of the corresponding edge. Let *P* = *x*_0_ *· · · x*_*m−*1_ be a stored path, and assume that the *i*- th node visit *x*_*i*_ corresponds to offset *j* in the list of path visits. We call (*x*_*i*_, *j*) the *GBWT position* for the visit. With the information stored in the record, we can compute efficiently the next GBWT position *LF* (*x*_*i*_, *j*) = (*x*_*i*+1_, *j*^*′*^). We use this ability in two ways. We can traverse a specific stored path from an arbitrary starting point. For any path *P* ^*′*^ we traverse in the graph, we can also maintain the interval [*a* … *b*) of path visits to the last handle *x*, where the corresponding prefixes of stored paths end with *P* ^*′*^. This allows us to determine the number of occurrences of any traversed path in the stored paths.

The other key component is the concatenation of node labels *l*(*v*) for all nodes *v ∈ V*. An index is used for finding the labels by node identifiers. We also store structured names (sample name, contig name, haplotype number, fragment number) for each original path. Tags (key–value pairs) can be used for storing additional metadata. Some vg tools assume that node labels are at most 1024 bp long. If the nodes of a graph have been chopped into smaller pieces for vg, GBZ can also store a translation between node identifiers in the chopped and the original graph.

### Stable graph names

Pangenome graphs built with Minigraph–Cactus (Hickey et al., 2024) come in several forms. The *full graph* built by aligning the assembly contigs is rarely used. We get the *default graph* by removing long unaligned segments such as centromeres from all haplotypes except the primary reference. By further removing rarely used non-reference nodes (e.g. those used by fewer than 10% of the haplotypes), we get the *frequency-filtered graph*, which is a better universal reference for read mapping and variant calling. We can also create *personalized graphs* to improve read mapping and variant calling accuracy by sampling local haplotypes from the default graph according to *k*-mer counts in the reads (Sirén et al., 2024).

By removing nodes and edges from a graph, we create a subgraph. If we have aligned reads to a subgraph, the supergraph is also a valid reference for the alignments. On the other hand, the indexes used in applications such as read mapping are specific to the graph, and they cannot be used with a subgraph or a supergraph. Further, while a graph with chopped nodes and the original graph represent the same alignment of the underlying haplotypes, alignments in one of them are not valid in the other without translating the coordinates. We therefore need a way of specifying which graphs can be used with which alignments or indexes.

With a linear reference sequence, we can use schemes such as refget (Yates et al., 2022; Campbell et al., 2025) for deriving a stable identifier by hashing the sequence itself. We have defined a similar scheme called *pggname* for pangenome graphs that hashes a minimal GFA representation of the graph in a canonical order with SHA-256. The pggname specification also defines a way of storing graph names and known subgraph and coordinate translation relationships as key–value tags and in GFA and GAF headers. See Supplement 3 for further details.

### Snarl decomposition

In applications such as variant calling, we expect that each component of a pangenome graph corresponding to linear chromosome has a linear high-level structure. Such a structure can be described using graph decompositions, such as the *snarl decomposition* (Paten et al., 2018) used in the vg toolkit.

Informally, a *snarl* is a subgraph separated by its two *boundary nodes* from the rest of the graph. Each graph component can be seen as a *top-level chain*, which is a sequence of nodes and snarls. The nodes represent shared sequence, while the snarls represent variation. When there is nested variation, the internal parts of a snarl (without its boundary nodes) can be seen as a set of disjoint chains and decomposed recursively.

Assume that we want to extract a subgraph corresponding to a reference interval. We first find the subpath representing that interval in the corresponding haplotype path. If both boundary nodes of a top-level snarl are in the subpath, the variation represented by the snarl is *contained* within the interval, and we should include the snarl in the subgraph. See Figure 2 for an example. We may also want to include *overlapping* snarls with at least one boundary node in the subpath. However, if the snarl is large, the subgraph may then end up representing a much longer reference interval than we intended.

**Figure 2.**
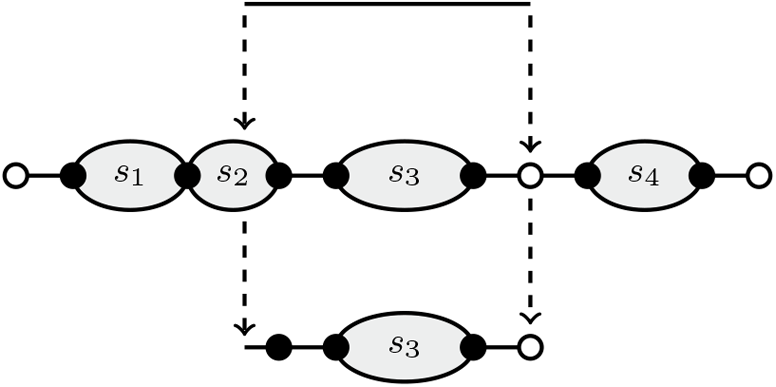
Middle: Ideal graph structure, with top-level snarls as gray ellipses, their boundary nodes as solid circles, and other nodes in the top-level chain as hollow circles. Top: A reference interval. Bottom: Subgraph corresponding to the reference interval with contained snarls. Extending the interval with overlapping snarls would also include snarl *s*_2_.

### Data compression

A typical data compression algorithm consists of three stages. In the *transformation* stage, we transform the data into another representation that we expect to be easier to compress. For example, Lempel–Ziv compressors such as *Zstandard* transform a string *S* over alphabet Σ into a sequence of substrings *S*[*i* … *i* + *l*) (defined by source position *i* and copy length *l*) copied from earlier in the string and explicit characters *c ∈* Σ. *Run-length encoding* transforms a sequence of *k* copies of character *c* into a pair (*c, k*).

In the *modeling* stage, we determine a probability distribution for each stream of *symbols*. Given an alphabet Σ, we assume that each symbol *c ∈* Σ occurs with probability *p*_*c*_ in the stream.

A Lempel–Ziv compressor may have separate streams for source positions, copy lengths, and explicit characters. The distribution may depend on the *context*. A zero-order model uses the same distribution for all symbols, while in an order-*k* model, the distribution depends on the *k* previous symbols.

The *encoding* stage transforms a sequence of symbols into a binary sequence according to the probability distribution.

Ideally the expected length of an encoded symbol should be close to the *entropy H* = *−* _*c∈*Σ_ *p*_*c*_ log_2_ *p*_*c*_ of the distribution. *Asymmetric numeral systems* (ANS) (Duda, 2014) are a computationally efficient way of achieving this. Variable-length integers are used when we assume that small numbers are more common than large ones, but determining or storing the exact distribution would be impractical. *Little-endian base 128* (LEB128) encodes an integer using 7 bits/byte in little-endian order and uses the high bit for determining whether the encoding continues in the next byte. It is often used when computational efficiency is more important than maximal compression.

File formats such as GAF consist of *records* with heterogeneous fields. If we compress all fields in a record together, we can decompress the record efficiently. *Columnar compression* handles each field across all records separately. It typically compresses the data better, at the expense of making decompression of entire records slower. *Block-based compression* is a common trade-off. It partitions the records into blocks and uses columnar compression within each block.

### GBZ-base

The GBZ file format was originally designed with memory-mapping in mind. This turned out to be too impractical. Instead of using memory-mapped GBZ files, we chose to design a new file format for interactive applications such as visualization. We chose an SQLite database as the container format and named this new file format *GBZ-base*.

Our main design goal was to preserve the data layout of GBZ, so that most of the data can be copied directly into the appropriate database tables. We merged GBWT node records and sequences into a single table and added smaller tables for structured path names and tags. Some GBZ components were left out. The structure used for identifying haplotype paths was already too inefficient in an in-memory graph, while the translation between the chopped graph and the original graph was rarely used. On the other hand, we added an index for translating positions in selected reference haplotypes to the corresponding handles.

### Node records in GBZ-base

A GBZ file has a node record for each handle. Sequence labels are stored only once for each node. In GBZ-base, we chose to merge the two in the `Nodes` table, with a record for each handle. This sequence duplication increases the size of a whole-genome human graph by approximately 1 GiB, which we consider an acceptable trade-off for convenience.

The Nodes table has the following fields:

- handle (integer): Handle *x* of the node visit; used as the primary key.
- edges (binary blob): List of outgoing edges using the same encoding as in GBWT.
- bwt (binary blob): List of path visits using the same encoding as in GBWT.
- sequence (binary blob): Encoded label *l*(*x*).
- next (integer): Optional link *next*(*x*) to the next handle in the chain.

The original GBZ used a custom sequence encoding, while version 2 switched to Zstandard. Both encodings compress the concatenated labels together. Because GBZ-base must be able to decode individual node records, we chose a simple encoding that packs three bases in a byte.

We use the `next` links for finding snarls. Let *x, y ∈ V* ^*′*^ be the visits to the boundary nodes of the snarl pointing towards the snarl. We then store the links *next*(*x*) *← y* and *next*(*y*) *← x*. If the result of a query contains handles *x* and *next*(*x*) and we want to extract a subgraph containing all variation covered by the result, we can extend the result with the subgraph between those two handles.

### Reference index

Table `ReferenceIndex` is used for mapping positions in the selected reference haplotypes to positions in the graph. It is functionally similar to the inverse suffix array samples used to support random access in an FM-index.

Let *i* be the integer identifier of an original path *P*_*i*_ we have selected for indexing, let *a* be a position in the underlying haplotype sequence corresponding to the start of handle *x* in the path, and let (*x, j*) be the corresponding GBWT position. We store sample (*i, a, x, j*) in `ReferenceIndex`, if *a* = 0 or the distance *a − a*^*′*^ from the previous sample (*i, a*^*′*^, *x*^*′*^, *j*^*′*^) is large enough (at least 1000 bp by default).

We use pairs (*i, a*) as the primary key of the table. Assume that we want to determine the graph position for sequence position *b* in path *P*_*i*_. We use the database index to find the last sampled position (*i, a, x, j*) such that *a ≤ b*. If *b − a <* |*l*(*x*)|, the graph position is sequence offset *b−a* in handle *x*. Otherwise we iterate with *a ← a* + |*l*(*x*)| and (*x, j*) *← LF* (*x, j*).

By default, we need almost 3 million samples taking tens of megabytes for each human haplotype. We can therefore afford indexing some haplotypes but not hundreds of them.

### Finding top-level chains

GBZ-base construction can import `next` links from top-level chains extracted from a snarl decomposition or a distance index (Chang et al., 2020) using `vg chains`. If precomputed toplevel chains are not provided, GBZ-base tries to find them using a simple algorithm. The algorithm only works in graph components with two tips (nodes with all edges on the same side) and a directed path between them. Minigraph–Cactus typically builds graphs like that.

The algorithm processes each weakly connected component separately. It first finds all articulation points in the component. Then it walks the shortest directed path between the tips and lists all articulation points that are visited exactly once. These are the nodes in the top-level chain for that component. Finally it determines which of them are boundary nodes of a snarl. Let *x ∈ V* ^*′*^ be a visit to a node *v ∈ V* in a top-level chain. Node *v* is a snarl boundary node and visit *x* points towards the snarl if and only if i) *x* has multiple successors, or ii) its only successor has multiple predecessors.

The semantics of the `next` links are the following: if handles *x* and *next*(*x*) are in the region of interest, the subgraph between them is also in the region. We store links between the boundary nodes of top-level snarls, as described earlier. If we have a unary path of two or more nodes between consecutive snarls, we also store links between the first and the last node in the path. This lets us traverse each top-level chain by following the `next` links. While the query mechanism also supports nested snarls, we currently do not store links between their boundary nodes.

### Subgraph queries in GBZ-base

Because the SQLite database engine is embedded in the user process, its query latency is lower than in client–server databases. It can answer a large number of small readonly queries (such as those retrieving individual node records) efficiently. However, the latency is still too high for traditional graph interfaces. We have therefore designed a query model based on extracting subgraphs around regions of interest into in-memory graph structures. See Supplement 4.1 for a more complete description of the query algorithms.

A query builds a subgraph in three steps. First we include the nodes covering the queried positions, which are typically expressed either as an interval in an indexed reference haplotype or as a set of node identifiers. Then we extend the subgraph to include a *greedy context* containing all nodes within a specified undirected distance (default 100 bp) of the queried positions. Finally, we may extend the subgraph with snarls. We can include only snarls that are contained in the subgraph or also those that are partially overlapping with it.

Once we have built the subgraph, we determine the local haplotype paths in it. If the query was based on a reference interval, we get proper metadata for the selected reference path, including the interval in the underlying haplotype sequence. We cannot identify the other haplotypes or determine their coordinates, as both GBZ-base and GBZ currently lack efficient data structures for that. Identical local haplotypes can optionally be merged. Query results can be processed in in-memory data structures or serialized in GFA or JSON formats.

### GAF-base

*GAF-base* stores sequence alignments to a pangenome graph in an SQLite database. It assumes a data model similar to the GAF specification used in vg (see Supplement 2). Target paths for the alignments are stored in a GBWT index. Other fields are stored using block-based compression similar to CRAM. Additional metadata can be stored as key–value tags.

### Node records in GAF-base

The GBWT index storing the target paths can be unidirectional (storing only the original paths) or bidirectional. It is stored in the `Nodes` table similar to the one in GBZ-base. The GAF-base version of the table does not include `next` links.

By default, we build a *reference-based* GAF-base without node labels. A reference-based GAF-base must be always used with a reference graph (a GBZ graph or a GBZ-base). We use the pggname scheme to list the appropriate reference graphs, if that information is available. We can also build a *reference-free* GAF-base by storing node labels. That increases the size of a whole-genome human database by approximately 2 GiB.

### Alignments

Each row in table `Alignments` stores information for a block of alignments. The default block size is 1000 alignments, which is appropriate for short reads. The blocks are indexed by (min handle, max handle) in the target paths. If all query sequences have the same length, that length is stored explicitly. The following binary blobs store the rest of the information:

- `gbwt starts` : GBWT position for the start of each path, with the handle stored relative to the minimum handle in the block. Encoded using LEB128.
- `names` : Concatenated query sequence / pair names compressed with Zstandard.
- `quality strings` : Concatenated base quality strings encoded with the ANS implementation from HTSlib (Bonfield, 2022). Currently rANS 4×16 with an order-1 model and run-length encoding.
- `difference strings` : Concatenated difference strings encoded using run-length encoding from GBWT. Inserted sequences use the same encoding as in the `Nodes` table.
- `flags` : Binary flags for each alignment (e.g. exact alignment, full-length alignment, properly paired).
- `numbers` : Numbers that cannot be derived from the other information (e.g. mapping quality, alignment score, aligned query / target intervals) encoded using LEB128.
- `optional` : Other optional fields concatenated and compressed with Zstandard.

### Building a GAF-base

GAF-base assumes that the alignments are sorted by (min handle, max handle) in the path. This can be done with the included `gafsort` tool or with `vg gamsort`. Both implement a multi-threaded multi-way external memory merge sort algorithm similar to GNU sort. The vg implementation is somewhat faster — likely due to a faster gzip decompressor. Default block size is 1 million alignments, which is appropriate for short reads. Merging is 32-way by default, allowing GAF files with up to 1024 blocks to be sorted in two rounds of merges.

The construction itself uses four threads: parser, encoder, insertion, and GBWT construction. The GBWT construction algorithm indexes the paths only in the forward orientation. It has been optimized for batches of short paths, all in the same graph region. It is therefore faster than the general-purpose algorithm used in vg. A pre-built GBWT index can also be provided to lower the memory usage of GAF-base construction.

### Subgraph queries in GAF-base

The primary GAF-base query extracts all alignments in a given subgraph. Since GAF is a reference-based format, this typically involves clipping each alignment into one or more fragments that are fully in the subgraph. See Supplement 4.2 for a more complete description of the query algorithms.

Let [*a* … *b*] be the minimal interval covering all handles in the subgraph, and let [*a*_*B*_ … *b*_*B*_] be the handle interval for block *B*. If the two intervals overlap, all alignments in the block are *candidates* we need to decompress. However, an overlap does not necessarily indicate that the alignments are in the subgraph. While pangenome graph construction tools try to assign similar identifiers to adjacent nodes, there are sometimes large gaps between nearby identifiers due to complex graph structures.

To mitigate this issue, we *cluster* the subgraph into multiple intervals [*a*_0_ … *b*_0_], [*a*_1_ … *b*_1_], … if there are large gaps between node identifiers. We then find the set of blocks overlapping with at least one of the intervals. Given a block of candidate alignments, we decompress the alignments and trace the target paths to determine if they are in the subgraph. Even with the mitigation, it often turns out that most candidates are not in the subgraph, if the subgraph is small.

## Results

We build GBZ-base for an HPRC human graph and GAF-bases for high-coverage human datasets. We then compare the size of a GAF-base to other sequence alignment formats and measure the performance of GBZ-base and GAF-base subgraph queries. See Supplement 8 for the specific commands used.

### Experimental setup

GBZ-base and GAF-base implementations are written in Rust. We measured their performance on two systems. *Laptop* is a 16” Apple MacBook Pro with an M2 Max (8 performance and 4 efficiency cores), 96 GiB RAM, and 2 TB SSD running macOS 26.5.1 (26.7 for subgraph queries). *Server* is an AWS r8id.16xlarge instance with 32 physical / 64 logical cores of Xeon 6975P-C, 512 GiB RAM, and 3.8 TB SSD running Ubuntu 24.04.4 LTS. We used the following software versions: GBZ-base 0.5.1 (0.6.2 for subgraph queries), vg 1.75.0 (Garrison et al., 2018), SAMtools 1.19.2 (Li et al., 2009), and KMC 3.2.4 (Kokot et al., 2017).

We used HPRC release 2 version 2.1 graphs built with Minigraph–Cactus using CHM13 as the primary reference (Lucas et al., 2026). See Supplement 5 for a discussion on using graphs built with the PanGenome Graph Builder (Garrison et al., 2024). The graphs were the evaluation versions, with samples HG002, HG005, and NA19240 left out, containing a total of 458 haplotypes. We aligned high-coverage Illumina NovaSeq, Element Biosciences, PacBio HiFi, and Oxford Nanopore R10 (ONT) reads for HG002 to the graphs using Giraffe. See Table 1 for details on the reads and Supplement 1 for data sources.

**Table 1.** Datasets used in GAF-base experiments. Number and total length of the reads in each dataset, and wall-clock time and peak memory usage for GAF sorting, GAF-base construction, and GAF-base decompression.

| Dataset | Reads | Total length | Sort | Build | Decompress |
| --- | --- | --- | --- | --- | --- |
| Element | 1,393,184,576 | 208.4 Gbp | 80 min, 2.5 GiB | 67 min, 65.0 GiB | 126 min, 9.4 GiB |
| Illumina | 948,769,000 | 143.2 Gbp | 27 min, 2.4 GiB | 39 min, 56.5 GiB | 68 min, 9.5 GiB |
| HiFi | 12,800,930 | 204.5 Gbp | 32 min, 12.2 GiB | 39 min, 37.6 GiB | 19 min, 9.2 GiB |
| ONT | 8,346,242 | 153.7 Gbp | 27 min, 14.4 GiB | 32 min, 37.8 GiB | 21 min, 9.2 GiB |

### GBZ-base construction

We built GBZ-base for the default graph. As it is a supergraph of the frequency-filtered graph and the base graph for personalized graphs, we can use it as a reference with reads aligned to those graphs. The construction took 451 seconds and 17.9 GiB memory on the laptop. About half of that time was used for finding top-level chains in the graph. The size of the database was 10.7 GiB, compared to 5.7 GiB for GBZ version 1 and 3.35 GiB for the experimental version 3.

**GAF-base construction**

We used the server for GAF-base construction. First we aligned the reads with Giraffe, using personalized graphs with short reads (Element, Illumina) and the frequency-filtered graph with long reads (HiFi, ONT) as the reference. Then we sorted the GAF output with the `gafsort` tool and built GAF-bases for the sorted alignments. We used blocks of 1 million alignments with short reads and 10 thousand alignments with long reads for sorting and blocks of 1000 alignments with short reads and 10 alignments with long reads in the database. See Table 1 for wall-clock times and peak memory usage.

Sorting took longer with Element reads than with the other datasets. Because we did 32-way merges, there were three rounds of merges for 1394 blocks. Noisy quality scores also made reading and writing compressed temporary files slower. GAF-base construction was more expensive with short reads, largely due to having to build a GBWT index with a very large number of very short paths. With Element reads, the database itself was also a bottleneck, as we had to insert much more data into the `Alignments` table due to noisy quality scores.

We also measured the time and memory usage for full decompression of the database. We used a naive algorithm that decompresses 100 blocks at a time using one thread and then passes them to another thread for serialization. See Table 1 for the results. This was reasonably fast with long reads but slower with short reads. The likely bottleneck with short reads was tracing the paths in the `Nodes` table. An optimized algorithm would read the paths into an in-memory GBWT index first.

### GAF-base size comparison

We compared the size of GAF-base to the sizes of the same alignments in other formats. See Table 2 for the results and Supplement 6 for the size breakdown of GAF-base components.

**Table 2.** File size comparison for the datasets in Table 1.

| Dataset | GAM | GAF | GAF (gz) | GAF (sorted, gz) | GAF-base | BAM | CRAM |
| --- | --- | --- | --- | --- | --- | --- | --- |
| Element | 226.8 GiB | 561.1 GiB | 117.4 GiB | 107.8 GiB | 90.2 GiB | 152.2 GiB | 75.8 GiB |
| Illumina | 113.4 GiB | 320.8 GiB | 42.6 GiB | 35.6 GiB | 27.4 GiB | 64.1 GiB | 18.0 GiB |
| HiFi | 116.6 GiB | 249.3 GiB | 32.3 GiB | 32.0 GiB | 22.3 GiB | 63.9 GiB | 18.2 GiB |
| ONT | 130.5 GiB | 195.4 GiB | 68.3 GiB | 68.1 GiB | 53.3 GiB | 91.6 GiB | 45.6 GiB |

GAM is the native alignment format used by the vg toolkit. GAF stores less information, but the files are larger due to the lack of built-in compression. Because GAF files are highly compressible, the initial output is typically compressed with (b)gzip. The size of compressed GAF decreases further when the alignments are sorted by handle intervals. GAF-base was between 16% and 30% smaller than sorted and compressed GAF. The BAM file in the results contains the same alignments projected to CHM13 with `vg surject`. The CRAM file was obtained by sorting the BAM and converting it to CRAM with SAMtools. In all cases, the size of GAF-base was between BAM and CRAM and closer to CRAM.

### Subgraph queries

We measured the performance of GBZ-base and GAF-base subgraph queries on the laptop. In each session, we queried 1000 random CHM13 intervals of a given length (100 bp, 1 kbp, 10 kbp, 100 kbp, or 1 Mbp). With 1 Mbp intervals, we limited the number of queries to 100 to avoid reading a too large fraction of the databases. We extracted a 100 bp greedy context around the query interval and contained snarls. We repeated each session with no reads, with Illumina reads, and with HiFi reads. Each session was started with a cold cache. We measured the wall-clock time required to read the subgraph and the alignments into in-memory data structures.

The results can be found in Figure 3. See Supplement 7 for the full results, including queries with a GBZ graph, with Element and ONT reads, and without snarls, as well as peak memory usage. With no reads, typical query times were in milliseconds for 100 bp and 1 kbp intervals, tens of milliseconds for 10 kbp intervals, hundreds of milliseconds for 100 kbp intervals, and seconds for 1 Mbp intervals. Queries in clipped regions were faster, as the subgraph consisted of a sequence of 1024 bp nodes. Queries for short intervals were faster with a GBZ graph, but loading it into memory and indexing the reference paths took 39 seconds and 12 GiB memory.

**Figure 3.**
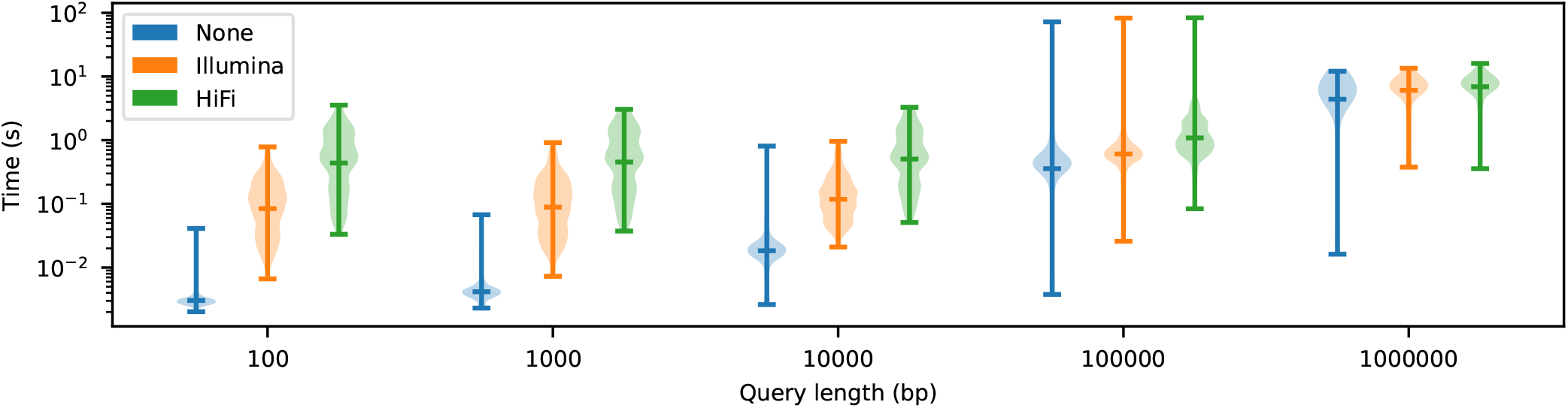
Violin plots for subgraph query times with random CHM13 intervals, 100 bp greedy context, and contained snarls. GBZ-base with no reads, Illumina reads, and HiFi reads.

If a non-reference haplotype has a large deletion within the greedy context, snarls may extend the reference interval beyond the intended length. In an extreme case, a 100 kbp query near 69.5 Mbp in chr7 was extended to over 10 Mbp, with the query taking 75 seconds. If we only include a greedy context or snarls but not both, we avoid these degenerate cases.

If we also query a GAF-base, typical query times start from tens or hundreds of milliseconds with 100 bp intervals and increase only a little with 10 kbp intervals. There are two main reasons for this behavior. First, read length becomes an effective context length around the subgraph. Second, when the subgraph is small, most candidate alignments do not overlap with it. See Supplement 7 for more details.

## Conclusion

GBZ-base is an indexed version of the GBZ file format. Due to database overhead, additional functionality, and storing the sequences in both orientations, GBZ-base files are typically almost twice as large as GBZ files. For best performance, the database should be stored on a local SSD. We can extract local subgraphs around given nodes or reference intervals in a fraction of a second, which makes the GBZ-base appropriate for interactive applications such as visualization. Context extension options should be used with care, to avoid accidentally extracting a much larger subgraph than intended.

GAF-base is an indexed binary format for sequence-to-graph alignments. It is compatible with the GAF format written by the vg toolkit. We can extract all alignments to a given local subgraph efficiently from a GAF-base. The performance of such queries depends on the size of the subgraph, the length of the reads, and the structure of the graph. We get the best performance when the subgraph can be defined by an interval of node identifiers.

While GAF-base files are smaller than other sequenceto-graph alignment formats, the GAF-base should not be understood as the pangenome equivalent of BAM or CRAM. The file format is still under active development. If mutability is not needed, the files can be made smaller by using a container format with less space overhead than SQLite. Compression can also be improved by following the behavior of CRAM more closely. For now, sorted (b)gzip-compressed GAF is still the best choice for long-term archival.

## Author contributions

J.S. conceived, designed and implemented GBZ-base and GAF-base. J.S. wrote the manuscript. B.P. consulted on the design and rationale for GBZ-base and GAF-base and edited the manuscript.

## Supplementary material

Supplementary material is available online.

## Data availability

No new data were generated or analyzed in support of this research.

## Human Pangenome Reference Consortium Authors

Derek Albracht^1^, Ivan A. Alexandrov^2^, Jamie Allen^3^, Alawi A. Alsheikh-Ali^4^, Nicolas Altemose^5^, Casey Andrews^6^, Dmitry Antipov^7^, Lucinda Antonacci-Fulton^1^, Mobin Asri^8^, Marcelo Ayllon^9^, Jennifer R. Balacco^10^, Floris P. Barthel^11^, Edward A. Belter Jr^1^, Halle D. Bender^8^, Andrew P. Blair^8^, Davide Bolognini^12^, Katherine E. Bonini^13^, Christina Boucher^14^, Guillaume Bourque^15,16,17^, Silvia Buonaiuto^18^, Shuo Cao^18^, Andrew Carroll^19^, Ann M. Mc Cartney^8^, Monika Cechova^8^, Mark J.P. Chaisson^20^, Pi-Chuan Chang^19^, Xian Chang^8^, Jitender Cheema^3^, Haoyu Cheng^21^, Claudio Ciofi^22^, Hiram Clawson^8^, Sarah Cody^1^, Vincenza Colonna^18^, Holland C. Conwell^23^, Robert Cook-Deegan^24^, Mark Diekhans^8^, Maria Angela Diroma^22^, Daniel Doerr^25,26,27^, Zheng Dong^6^, Danilo Dubocanin^5^, Richard Durbin^28,29^, Jana Ebler^25,30^, Evan E. Eichler^9,31^, Jordan M. Eizenga^8^, Parsa Eskandar^8^, Eddie Ferro^14^, Anna-Sophie Fiston-Lavier^32,33^, Sarah M. Ford^23^, Willard W. Ford^34^, Giulio Formenti^10^, Adam Frankish^3^, Mallory A. Freeberg^3^, Qichen Fu^6^, Stephanie M. Fullerton^35^, Robert S. Fulton^1^, Shenghan Gao^36^, Yan Gao^37^, Gage H. Garcia^9^, Obed A. Garcia^38^, Joshua M.V. Gardner^8^, Shilpa Garg^39^, Erik Garrison^18^, Nanibaa’ A. Garrison^40,41,42^, John E. Garza^1^, Margarita Geleta^43^, Mohammadmersad Ghorbani^44^, Tina A. Graves-Lindsay^1^, Richard E. Green^23^, Carol W. Greider^45^, Cristian Groza^46^, Bida Gu^20^, Andrea Guarracino^11,18^, Melissa Gymrek^47^, Maximilian Haeussler^8^, Leanne Haggerty^3^, Ira M. Hall^48,49^, Nancy F. Hansen^7^, Yue Hao^11^, Mohammad Amiruddin Hashmi^4^, David Haussler^8^, Prajna Hebbar^8^, Peter Heringer^25,26,27^, Glenn Hickey^8^, Todd L. Hillaker^8^, S. Nakib Hossain^3^, Neng Huang^37,50^, Sarah E. Hunt^3^, Toby Hunt^3^, Alexander G. Ioannidis^5,8^, Nafiseh Jafarzadeh^8^, Nivesh Jain^10^, Erich D. Jarvis^10,31^, Maryam Jehangir^11^, Juan Jiang^6^, Eimear E. Kenny^13^, Juhyun Kim^7^, Bonhwang Koo^10^, Sergey Koren^7^, Milinn Kremitzki^1,6^, Charles H. Langley^51^, Ben Langmead^52^, Heather A. Lawson^6^, Daofeng Li^6^, Heng Li^37,50^, Wen-Wei Liao^48,49^, Jiadong Lin^9^, Tianjie Liu^6^, Glennis A. Logsdon^36^, Ryan Lorig-Roach^8^, Jonathan LoTempio Jr^53^, Hailey Loucks^8^, Jane E. Loveland^3^, Jianguo Lu^54^, Shuangjia Lu^48,49^, Julian K. Lucas^8^, Walfred Ma^20^, Juan F. Macias-Velasco^1,6,55^, Kateryna D. Makova^56^, Maximillian G. Marin^37,50^, Christopher Markovic^1^, Tobias Marschall^25,30^, Franco L. Marsico^18^, Fergal J. Martin^3^, Mira Mastoras^8^, Capucine Mayoud^32^, Brandy McNulty^8^, Jack A. Medico^10^, Julian M. Menendez^8^, Karen H. Miga^8^, Anna Minkina^57^, Matthew W. Mitchell^58^, Saswat K. Mohanty^59^, Younes Mokrab^44,60,61^, Jean Monlong^62^, Shabir Moosa^44^, Avelina Moreno-Ochando^63,64^, Shinichi Morishita^65^, Jonathan M. Mudge^3^, Katherine M. Munson^9^, Njagi Mwaniki^66^, Nasna Nassir^4^, Chiara Natali^22^, Shloka Negi^8^, Lingbin Ni^9^, Adam M. Novak^8^, Faith Okamoto^8^, Keisuke K. Oshima^36^, Pilar N. Ossorio^67^, Chie Owa^65^, Sadye Paez^10^, Benedict Paten^8^, Clelia Peano^12,68^, Adam M. Phillippy^7^, Brandon D. Pickett^7^, Laura Pignata^18^, Nadia Pisanti^66^, David Porubsky^9,69^, Pjotr Prins^18^, Timofey Prodanov^25,30^, Anandi Radhakrishnan^8^, T. Rhyker Ranallo-Benavidez^11^, Brian J. Raney^8^, Mikko Rautiainen^70^, Alessandro Raveane^12^, Andreas Rechtsteiner^45^, Luyao Ren^9,31^, Arang Rhie^7^, Fedor Ryabov^71,72^, Samuel Sacco^23^, Farnaz Salehi^18^, Michael C. Schatz^52,73^, Laura B. Scheinfeldt^74^, Aarushi Sehgal^34^, William E. Seligmann^23^, Mahsa Shabani^75^, Kishwar Shafin^19^, Shadi Shahatit^32^, Ruhollah Shemirani^13^, Vikram S. Shivakumar^52^, Swati Sinha^3^, Jouni Sirén^8^, Linnéa Smeds^59^, Steven J. Solar^7^, Marco Sollitto^10,22^, Nicole Soranzo^12,28,76^, Andrew B. Stergachis^9,57^, Marie-Marthe Suner^3^, Yoshihiko Suzuki^65^, Arda Söylev^25,30^, Ahmad Abou Tayoun^77,78^, Jack A.S. Tierney^3^, Chad Tomlinson^1^, Francesca Floriana Tricomi^3^, Mohammed Uddin^4,79^, Matteo Tommaso Ungaro^23,80^, Rahul Varki^14^, Flavia Villani^18^, Ivo Violich^8^, Mitchell R. Vollger^57^, Brian P. Walenz^7^, Charles Wang^81^, Lisa E. Wang^13^, Ting Wang^1,6,55^, Aaron M. Wenger^82^, Conor V. Whelan^10^, Zilan Xin^6^, Zheng Xu^6^, Kai Ye^83^, DongAhn Yoo^9^, Wenjin Zhang^6^, Ying Zhou^37^, Xiaoyu Zhuo^6^, Giulia Zunino^12^

^1^ McDonnell Genome Institute, Washington University School of Medicine, St. Louis, MO 63108, USA

^2^ Department of Human Molecular Genetics and Biochemistry, Faculty of Medical and Health Sciences, Tel Aviv University, Tel Aviv 69978, Israel

^3^ European Molecular Biology Laboratory, European Bioinformatics Institute (EMBL-EBI), Wellcome Genome Campus, Hinxton, Cambridge CB10 1SD, UK

^4^ Center for Applied and Translational Genomics (CATG), Mohammed Bin Rashid University of Medicine and Health Sciences, Dubai Health, Dubai, UAE

^5^ Department of Genetics, Stanford University, Palo Alto, CA 94304 USA

^6^ Department of Genetics, Washington University School of Medicine, St. Louis, MO 63110, USA

^7^ Genome Informatics Section, Center for Genomics and Data Science Research, National Human Genome Research Institute, National Institutes of Health, Bethesda, MD 20892, USA

^8^ UC Santa Cruz Genomics Institute, University of California, Santa Cruz, CA 95060, USA

^9^ Department of Genome Sciences, University of Washington School of Medicine, Seattle, WA 98195, USA

^10^ The Vertebrate Genome Laboratory, The Rockefeller University, New York, NY 10065, USA

^11^ Bioinnovation and Genome Sciences, The Translational Genomics Research Institute (TGen), Phoenix, AZ 85004, USA

^12^ Human Technopole, Milan, Italy

^13^ Institute for Genomic Health, Icahn School of Medicine at Mount Sinai, New York, NY 10029, USA

^14^ Department of Computer and Information Science and Engineering, University of Florida, Gainesville, FL 32611, USA

^15^ Canadian Center for Computational Genomics, McGill University, Montréal, QC H3A 0G1, Canada

^16^ Department of Human Genetics, McGill University, Montréal, QC H3A 0G1, Canada

^17^ Victor Phillip Dahdaleh Institute of Genomic Medicine, Montréal, QC H3A 0G1, Canada

^18^ Department of Genetics, Genomics and Informatics, University of Tennessee Health Science Center, Memphis, TN 38163, USA

^19^ Google LLC, Mountain View, CA 94043, USA

^20^ Quantitative and Computational Biology, University of Southern California, Los Angeles, CA 90089, USA

^21^ Department of Biomedical Informatics and Data Science, Yale School of Medicine, New Haven, CT 06510, USA

^22^ Department of Biology, University of Florence, Sesto Fiorentino, FI 50019, Italy

^23^ Department of Ecology and Evolutionary Biology, University of California, Santa Cruz, CA 95060, USA

^24^ Arizona State University, Consortium for Science, Policy & Outcomes, Washington, DC 20006, USA

^25^ Center for Digital Medicine, Heinrich Heine University Düsseldorf, Düsseldorf, NRW, DE

^26^ Department for Endocrinology and Diabetology at the Medical Faculty and University Hospital Düsseldorf, Heinrich Heine University Düsseldorf, Düsseldorf, NRW, DE

^27^ Paul-Langerhans-Group Computational Diabetology, German Diabetes Center (DDZ) and Leibniz Institute for Diabetes Research, Düsseldorf, NRW, DE

^28^ Wellcome Sanger Institute, Genome Campus, Hinxton, CB10 1RQ, UK

^29^ Department of Genetics, University of Cambridge, Cambridge, CB2 3EH, UK

^30^ Institute for Medical Biometry and Bioinformatics, Medical Faculty and University Hospital Düsseldorf, Heinrich Heine University, Düsseldorf, NRW, DE

^31^ Howard Hughes Medical Institute, Chevy Chase, MD 20815, USA

^32^ ISEM, Univ Montpellier, CNRS, IRD, Montpellier, FR

^33^ Institut Universitaire de France, Paris, FR

^34^ Department of Computer Science and Engineering, University of California San Diego, La Jolla, CA 92093, USA

^35^ Department of Bioethics & Humanities, University of Washington School of Medicine, Seattle, WA 98195, USA

^36^ Department of Genetics, Epigenetics Institute, Perelman School of Medicine, University of Pennsylvania, Philadelphia, PA 19104, USA

^37^ Department of Data Science, Dana-Farber Cancer Institute, Boston, MA 02215, USA

^38^ Department of Anthropology, University of Kansas, Lawrence, KS 66045, USA

^39^ School of Health Sciences, University of Manchester, Manchester M13 9PL, UK

^40^ Traditional, ancestral and unceded territory of the Gabrielino/Tongva peoples, Institute for Society & Genetics, University of California, Los Angeles, Los Angeles, CA 90095, USA

^41^ Traditional, ancestral and unceded territory of the Gabrielino/Tongva peoples, Institute for Precision Health, David Geffen School of Medicine, University of California, Los Angeles, Los Angeles, CA 90095, USA

^42^ Traditional, ancestral and unceded territory of the Gabrielino/Tongva peoples, Division of General Internal Medicine & Health Services Research, David Geffen School of Medicine, University of California, Los Angeles, Los Angeles, CA 90095, USA

^43^ Department of Electrical Engineering and Computer Science, University of California, Berkeley, Berkeley, CA 94720, USA

^44^ Medical and Population Genomics Lab, Sidra Medicine, Doha, Qatar

^45^ Department of Molecular Cell and Developmental Biology, University of California, Santa Cruz, CA, USA

^46^ Montreal Heart Institute, Montréal, QC, Canada

^47^ Department of Pediatrics, University of California San Diego, La Jolla, CA 92093, USA

^48^ Center for Genomic Health, Yale University School of Medicine, New Haven, CT 06510, USA

^49^ Department of Genetics, Yale University School of Medicine, New Haven, CT 06510, USA

^50^ Department of Biomedical Informatics, Harvard Medical School, Boston, MA 02115, USA

^51^ Department of Evolution and Ecology and the Center for Population Biology, University of California, One Shields, Davis, CA 95616, USA

^52^ Department of Computer Science, Johns Hopkins University, Baltimore, MD 21218, USA

^53^ Department of Pediatrics, Division of Genetics, School of Medicine, University of California, Irvine, CA 92697, USA

^54^ Sun Yat-sen University, Guangzhou, China

^55^ Edison Family Center for Genome Sciences & Systems Biology, Washington University School of Medicine, St. Louis, MO 63110, USA

^56^ Department of Biology and Center for Medical Genomics, Penn State University, University Park, PA 16802, USA

^57^ Division of Medical Genetics, Department of Medicine, University of Washington School of Medicine, Seattle, WA 98195, USA

^58^ The Jackson Laboratory for Genomic Medicine, Farmington, CT 06032, USA

^59^ Department of Biology, Penn State University, University Park, PA 16802, USA

^60^ Department of Biomedical Science, College of Health Sciences, Qatar University, Doha, Qatar

^61^ Department of Genetic Medicine, Weill Cornell Medicine-Qatar, Doha, Qatar

^62^ IRSD - Digestive Health Research Institute, University of Toulouse, INSERM, INRAE, ENVT, UPS, Toulouse, FR

^63^ MATCH biosystems, S.L., Elche, Spain

^64^ Universidad Miguel Hernández de Elche, Elche, Spain

^65^ Department of Computational Biology and Medical Sciences, The University of Tokyo, Kashiwa, Chiba 277-8561, Japan

^66^ Department of Computer Science, University of Pisa, Pisa, Italy

^67^ Law School, University of Wisconsin-Madison, Madison, WI 53706, USA

^68^ Institute of Genetics and Biomedical Research, UoS of Milan, National Research Council, Milan, Italy

^69^ Genome Biology Unit, European Molecular Biology Laboratory (EMBL), Heidelberg, DE

^70^ Institute for Molecular Medicine Finland, Helsinki Institute of Life Science, University of Helsinki, Helsinki, Finland

^71^ The Center for Bio- and Medical Technologies, Moscow, RUS

^72^ Centre for Biomedical Research and Technology, HSE University, Moscow, RUS

^73^ Department of Biology, Johns Hopkins University, Baltimore, MD 21218, USA

^74^ Coriell Institute for Medical Research, Camden, NJ 08103, USA

^75^ University of Amsterdam, Amsterdam, Netherlands

^76^ School of Clinical Medicine, University of Cambridge, Cambridge, CB2 0SP, UK

^77^ Center for Genomic Discovery, Mohammed Bin Rashid University, Dubai Health, UAE

^78^ Dubai Health Genomic Medicine Center, Dubai Health, UAE

^79^ GenomeArc Inc, Mississauga, ON, Canada

^80^ Department of Biology and Biotechnologies “Charles Darwin”, University of Rome “La Sapienza”, Rome 00185, IT

^81^ Center for Genomics, Loma Linda University School of Medicine, Loma Linda, CA 92350, USA

^82^ PacBio, Menlo Park, CA 94025, USA

^83^ The first affiliated hospital of Xi’an Jiaotong University, Xi’an Jiaotong University, Xi’an, Shaanxi, 710049, China

