## Supplementary Material for "GBZ-base and GAF-base: Indexed pangenome file formats"

September 28, 2026

#### 1 Data sources

##### 1.1 Graphs

We used HPRC release 2 version 2.1 evaluation graphs with CHM13 as the primary reference from https://s3-us-west-2.amazonaws.com/human-pangenomics/index.html?prefix=pangenomes/freeze/release2/minigraph-cactus/v2.1/benchmark-graphs/hprc-v2.1-mc-chm13-eval/. These graphs do not contain samples HG002, HG005, and NA19240.

- Default graph: hprc-v2.1-mc-chm13-eval.gbz (`default.gbz` in the scripts)
- Haplotype index for the default graph: hprc-v2.1-mc-chm13-eval.hapl (`default.hapl`)
- Frequency-filtered graph: hprc-v2.1-mc-chm13-eval.d46.gbz (`filtered.gbz`)

We also used the HPRC release 2 whole-genome PGGB graph: https://s3-us-west-2.amazonaws.com/human-pangenomics/pangenomes/freeze/release2/pggb/gfas/whole-genome/20250930_hprc25272.p98-k311.tmp.fix.gfa.zst.

##### 1.2 Reads

We reused the high-coverage reads for HG002 from the long read Giraffe paper [2]. The reads are available at: https://cgl.gi.ucsc.edu/data/lr-giraffe/reads/real/HG002/.

- Element Biosciences: HG002.GAT-LI-C044.fastq.gz (`element.fq.gz`). Originally from the Telomere-to-Telomere Consortium [3].
- Illumina NovaSeq: HG002.novaseq.pcr-free.40x.fq.gz (`illumina.fq.gz`). Originally from Google's gold-standard benchmarking dataset collection [1].
- PacBio HiFi: HG002Revio_hg002v1.0.1_hifi_revio_pbmay24.pri.unshuffled.fastq.gz (`hifi.fq.gz`). Originally from the Telomere-to-Telomere Consortium [3].
- ONT R10: r10y2025.HG002_PAW70337.fastq.gz (`ont.fq.gz`). Originally from the Genome in a Bottle data release 2025.01 by Oxford Nanopore Technologies [6].

### 2 The Graph Alignment Format (GAF)

The latest version of this document is hosted at:
https://github.com/vgteam/vg/blob/master/doc/static/GAF.md

This document describes version 1.0 of the vg interpretation of the Graph Alignment Format (GAF). It is a superset of a subset of the original GAF format. That format in turn is a superset of the PAF format. Sequence names and optional fields follow conventions set in the SAM format. Difference strings are defined in the minimap2 man page. Paths are represented as GFA walks. Reference graphs may have pggname stable names.

#### 2.1 Overview

GAF is a tab-delimited file format for sequence alignments to bidirected sequence graphs. The file is encoded in UTF-8. Unless otherwise specified, all fields are restricted to 7-bit US-ASCII.

Each file consists of a number of header lines followed by a number of alignment lines. Each line can be split into a number of fields separated by TAB (`\t`) characters.

#### 2.2 Typed fields

Typed fields are stored in the SAM-style `TAG:TYPE:VALUE` format. The tag is a two-character string matching `[A-Za-z][A-Za-z0-9]`.

The following types are currently supported:

| Type | Description |
|---|---|
| `A` | Printable character in `[!-~]` |
| `Z` | String of printable characters and spaces (`[ !-~]*`) |
| i | Signed 64-bit integer |
| f | Double-precision floating point number |
| b | Boolean value, with 1 for true and 0 for false |

#### 2.3 Header lines

##### Since vg 1.70.0

Header lines are optional, and they must all appear before the first alignment line. The first field of each header line is a three-character tag matching `@[A-Za-z][A-Za-z0-9]`.

**Example:**

```
@HD VN:Z:1.0
@RN 7f4b28c71ceb808aebd8b8e9fe85e79d0d208ee263ffe9fcdef5ade20534ceb5
@SG 7f4b28c71ceb808aebd8b8e9fe85e79d0d208ee263ffe9fcdef5ade20534ceb5
    e10f3b362d8a4273059d9aea38a78bd71913418c3f3c9a2b5ea44e86de2c1181
@TL e10f3b362d8a4273059d9aea38a78bd71913418c3f3c9a2b5ea44e86de2c1181
    1f133f116e8dd98fc07a647a8954038c2bcf07a45759ba94718471fe34ed7a7c
```

###### 2.3.1 File headers

File headers start with tag `@HD`. They may contain any number of optional typed fields. The following optional fields are known.

| Tag | Type | Description |
|---|---|---|
| `VN` | `Z` | Version number (e.g. 1.0; only one allowed in the file) |

###### 2.3.2 Reference name

**Since vg 1.71.0**

The graph the sequences were aligned to can be identified using a reference name line. A reference name line starts with tag `@RN` and contains the pggname (SHA-256 hash of the canonical GFA representation) of the graph as the second field. There may be optional typed fields. There can be only one reference name line in a file.

###### 2.3.3 Graph relationships

**Since vg 1.71.0**

If the sequences were aligned to graph A, which is a subgraph of B, graph B is also a valid reference for the alignments. If there is a known coordinate translation from graph B to graph C, graph C can also be used as a reference after translating the coordinates. Subgraph (`@SG`) and translation (`@TL`) lines can used to describe such relationships between reference graphs.

A subgraph line contains the pggname of the subgraph as the second field and the name of the supergraph as the third field. A translation line contains the name of the source graph as the second field and the name of the destination graph as the third field. Both line types may contain optional typed fields, and there may be any number of such lines.

##### 2.4 Alignment lines

Each alignment line has 12 mandatory fields. Missing values in fields 3 to 11 are indicated by character `*`.

| Field | Type | Description |
|---|---|---|
| 1 | `Z` | Query sequence name |
| 2 | `i` | Query sequence length |
| 3 | `i` | Query start (0-based; closed) |
| 4 | `i` | Query end (0-based; open) |
| 5 | `A` | Strand relative to the path; always `+` |
| 6 | `Z` | Target path represented as a GFA walk |
| 7 | `i` | Target path length |
| 8 | `i` | Start position on the target path (0-based; closed) |
| 9 | `i` | End position on the target path (0-based; open) |
| 10 | `i` | Number of matches |
| 11 | `i` | Number of matches, mismatches, insertions, and deletions |
| 12 | `i` | Mapping quality (0-255; 255 for missing) |

**Example:**

```
read1   6   0   6   +   >2>3>4  12  2   8   6   6   60  cs:Z::6
read2   7   0   7   +   >2>5>6  11  1   8   7   7   60  cs:Z::7
read3   7   0   7   *   *   *   *   *   *   *   255 cs:Z:+GATTACA
```

###### 2.4.1 Query sequence name

Query sequence names must follow SAM conventions. A name may contain any printable ASCII characters in the range `[!-~]`, except `@`. This allows distinguishing header lines from alignment lines.

###### 2.4.2 Target path

This version of GAF does not allow specifying the target path using stable rGFA coordinates or nodes (GFA segments) with string names. Nodes must have positive integer identifiers. Node identifier `0` cannot be used, as many graph implementations reserve it for technical purposes.

###### 2.4.3 Optional fields

Optional fields are SAM-style typed fields. No tag can appear more than once on the same line, and the order of the optional fields does not matter.

###### 2.4.4 Difference string

Difference strings represent an edit script that transforms the given interval of the target path to the given interval of the query sequence. They are stored as an optional field `cs` of type `Z`. We support a subset of the operations defined for minimap2 difference strings.

| **Operation** | **Regex** | **Description** |
|---|---|---|
| : | `[0-9]+` | Number of matching bases |
| * | `[ACGTN][ACGTN]` | Mismatch as (target base, query base) |
| + | `[ACGTN]+` | Insertion as the unaligned query bases |
| - | `[ACGTN]+` | Deletion as the unaligned target bases |

###### 2.4.5 Other defined optional fields

| **Tag** | **Type** | **Description** |
|---|---|---|
| AS | i | Alignment score |
| bq | Z | Base quality string; must have the same length as the query sequence |
| fn | Z | Name of the next fragment (for paired alignments; cannot be used with `fp`) |
| fp | Z | Name of the previous fragment (for paired alignments; cannot be used with `fn`) |
| pd | b | This alignment and its pair (specified by `fn` or `fp`) are properly paired |
| fi | i | Fragment identifier for a fragmented alignment (see below) |

##### 2.5 Conventions

###### 2.5.1 Header lines

**Since vg 1.70.0**

The first line of a GAF file is a file header (`@HD`) with the version number (`VZ:Z`) tag. Any additional file header lines follow. Other types of header lines are after file header lines.

###### 2.5.2 Primary alignments

A primary alignment represents an alignment of the entire query sequence to a non-empty interval of a target path. Query start (field 3) must be `0` and query end (field 4) must have the same value as query sequence length (field 2). A difference string must be present to allow recovering the entire query sequence.

###### 2.5.3 Unaligned sequences

**Since vg 1.70.0**

An unaligned sequence is represented as an alignment of the entire query sequence to a missing interval of a missing target path. Query start (field 3) must be `0` and query end (field 4) must have the same value as query sequence length (field 2). A difference string must be present, with the entire query sequence as a single insertion, to allow recovering the sequence.

###### 2.5.4 Fragmented alignments

A fragmented alignment is a single alignment represented as number of alignment lines (e.g. corresponding to subpaths that are within a specific subgraph). The fragments (alignment lines) correspond to non-overlapping intervals of the underlying alignment. Each fragment represents an alignment of a non-empty query interval to a non-empty interval of a non-empty target path.

Query interval (fields 3 and 4), target path (fields 6 to 9), and the difference string must be specific to each fragment. Alignment statistics (fields 10 to 12) may be inherited from the underlying alignment or be specific to each fragment. Fragments are identified by fragment indexes starting from `1`, stored as an optional field `fi` of type `i`.

#### 3 Stable names for pangenome graphs

The latest version of this document is hosted at:
https://github.com/jltsiren/pggname/blob/main/README.md

This is a proposal for generating stable names for pangenome graphs. The names are SHA-256 hashes of a canonical GFA representation of the graph.

See refget for a similar naming scheme for sequences.

#### 3.1 Intended applications

- Tagging various indexes with the name of the corresponding graph.
- As a reference name in a read alignment file.
- For representing relationships such as ”A is a subgraph of B” or ”A can be translated to B”.
  - If A is a subgraph of B, graph B can be used as a reference with reads aligned to A.
  - Some tools chop long nodes to smaller fragments, but coordinates in the chopped graph can be translated to the original coordinates.

#### 3.2 Example

We have three graphs:

- `original.gfa`: The original graph with some long nodes.
- `translated.gbz`: The same graph, with long nodes chopped into 1024 bp fragments.
- `sampled.gbz`: A personalized graph sampled from `translated.gbz`.

These graphs have the following names:

```
1f133f116e8dd98fc07a647a8954038c2bcf07a45759ba94718471fe34ed7a7c
    original.gfa
e10f3b362d8a4273059d9aea38a78bd71913418c3f3c9a2b5ea44e86de2c1181
    translated.gbz
7f4b28c71ceb808aebd8b8e9fe85e79d0d208ee263ffe9fcdef5ade20534ceb5
    sampled.gbz
```

We want to store the following information for `sampled.gbz`:

- The name of the graph.
- `sampled.gbz` is a subgraph of `translated.gbz`.
- Coordinates can be translated in both directions between `translated.gbz` and `original.gfa`.

##### 3.2.1 GBZ tags

```
pggname = 7f4b28c71ceb808aebd8b8e9fe85e79d0d208ee263ffe9fcdef5ade20534ceb5
subgraph = 7f4b28c71ceb808aebd8b8e9fe85e79d0d208ee263ffe9fcdef5ade20534ceb5,
    e10f3b362d8a4273059d9aea38a78bd71913418c3f3c9a2b5ea44e86de2c1181
translation = e10f3b362d8a4273059d9aea38a78bd71913418c3f3c9a2b5ea44e86de2c1181,
    1f133f116e8dd98fc07a647a8954038c2bcf07a45759ba94718471fe34ed7a7c;
    1f133f116e8dd98fc07a647a8954038c2bcf07a45759ba94718471fe34ed7a7c,
    e10f3b362d8a4273059d9aea38a78bd71913418c3f3c9a2b5ea44e86de2c1181
```

##### 3.2.2 GFA header

```
H   NM:Z:7f4b28c71ceb808aebd8b8e9fe85e79d0d208ee263ffe9fcdef5ade20534ceb5
H   SG:Z:7f4b28c71ceb808aebd8b8e9fe85e79d0d208ee263ffe9fcdef5ade20534ceb5,
    e10f3b362d8a4273059d9aea38a78bd71913418c3f3c9a2b5ea44e86de2c1181
H   TL:Z:e10f3b362d8a4273059d9aea38a78bd71913418c3f3c9a2b5ea44e86de2c1181,
    1f133f116e8dd98fc07a647a8954038c2bcf07a45759ba94718471fe34ed7a7c
H   TL:Z:1f133f116e8dd98fc07a647a8954038c2bcf07a45759ba94718471fe34ed7a7c,
    e10f3b362d8a4273059d9aea38a78bd71913418c3f3c9a2b5ea44e86de2c1181
```

##### 3.2.3 GAF header

```
@RN 7f4b28c71ceb808aebd8b8e9fe85e79d0d208ee263ffe9fcdef5ade20534ceb5
@SG 7f4b28c71ceb808aebd8b8e9fe85e79d0d208ee263ffe9fcdef5ade20534ceb5
    e10f3b362d8a4273059d9aea38a78bd71913418c3f3c9a2b5ea44e86de2c1181
@TL e10f3b362d8a4273059d9aea38a78bd71913418c3f3c9a2b5ea44e86de2c1181
    1f133f116e8dd98fc07a647a8954038c2bcf07a45759ba94718471fe34ed7a7c
@TL 1f133f116e8dd98fc07a647a8954038c2bcf07a45759ba94718471fe34ed7a7c
    e10f3b362d8a4273059d9aea38a78bd71913418c3f3c9a2b5ea44e86de2c1181
```

Here we use `RN` (reference name) instead of `NM` (name).

#### 3.3 Canonical GFA format

Sort the nodes by their identifiers. Interpret node identifiers as integers, if possible, and fall back to strings if at least one of the identifiers is not an integer.

For each node, in sorted order, output:

- S-line for the node without optional fields.
- L-lines for all canonical edges, without the overlap field or optional fields, in sorted order.

The canonical GFA representation of the graph does not include any other information, such as header lines, paths, or walks.

##### 3.3.1 Technicalities

Each line is terminated by a single `\n`, and the fields in a line are separated by a single `\t`. There are no empty fields or empty lines. The content of each field must be as in valid GFA.

##### 3.3.2 Nodes (GFA segments)

The sequence label of each node must be stored explicitly. Sequences are case sensitive, as some graph implementations do not normalize them. Upper case sequences are strongly recommended.

###### 3.3.3 Edges (GFA links)

An edge is canonical, if the source id is smaller than the destination id. A self-loop is canonical, if at least one of the nodes is in forward orientation.

Edges are sorted by (source orientation, destination id, destination orientation). The forward orientation comes before the reverse orientation.

###### 3.3.4 Example

Consider the following example graph from the GFA specification, with overlaps changed to `0M`:

```
H   VN:Z:1.0
S   11  ACCTT
S   12  TCAAGG
S   13  CTTGATT
L   11  +   12  -   0M
L   12  -   13  +   0M
L   11  +   13  +   0M
P   14  11+,12-,13+ 0M,0M
```

Its canonical GFA representation is:

```
S   11  ACCTT
L   11  +   12  -
L   11  +   13  +
S   12  TCAAGG
L   12  -   13  +
S   13  CTTGATT
```

And its stable name is:

```
54b49d18354a34fbd1af9aaac279e1b3ee67b2f68f0ff79f5ebf6c50c8d922a5
```

#### 4 Subgraph query algorithms

This section describes the algorithms and data structures used for subgraph queries, as they are implemented in GBZ-base version 0.6.2.

##### 4.1 Querying a GBZ-base or a GBZ graph

A subgraph query extracts a subgraph from a GBZ-base or a GBZ graph into an in-memory data structure. The subgraph data structure consists of a B-tree mapping handles to node records and a list of paths. Node records are the same as in GBZ-base, except that the edges are decompressed into an array of (destination handle, offset) pairs. Each path is a list of handles. There may also be optional metadata for the subgraph and the paths.

##### 4.1.1 Finding reference positions

If the query is based on a reference position, we first need to find the node covering the query. (For queries based on reference intervals, we find the node covering the start of the interval.) The first step towards that is finding the identifier and metadata for the path that covers the query. This is currently done with a linear scan over path names, as we expect that there are no more than tens of thousands of paths. Future versions may add an index for path names.

Next we need to find the last indexed position at or before the query position. With a GBZ-base, we query table `ReferenceIndex`, as described in the main text. That query is effectively a predecessor query in a database index. With a GBZ graph, we use an in-memory index we built after loading the graph. The index uses Elias–Fano encoded bitvectors [5] that mark the indexed positions in the reference paths.

Finally we need to follow the path forward to the query position, as described in the main text. For each node we encounter, we tentatively add the records for both orientations of the node to the subgraph. Then we use the information stored in the extracted records to compute LF-mapping to find the next node in the path.

##### 4.1.2 Adding greedy context

Context extraction is based on undirected shortest distances in a graph, where the left and right sides of a node in the pangenome graph are considered separate nodes. The distance between node sides connected by an edge in the pangenome graph is always 1, while the other side of node $v$ can be reached by crossing distance $|\ell(v)| - 1$ over the node. In some cases, the shortest path to the other side does not cross the node itself.

We use Dijkstra's algorithm, with active node sides in a binary heap, visited node sides in a B-tree, and the identifiers of the nodes to be removed in a B-tree. The last one is initialized with the nodes currently stored in the subgraph structure, as we have cached them but do not know if they are actually in the subgraph. When we visit a node side, we remove it from the active sides and add it to the visited sides. We also remove the node from those to be removed. If the node is not already in the subgraph, we add the records in both orientations. After we have visited all node sides within the context length, we go through the nodes to be removed and remove them from the subgraph.

If the query is based on a reference position, we initialize active node sides with both sides of the node covering the position, with appropriate distances to them. For queries based on a set of nodes, we start from both sides of each node, with distance 0 to them. With queries based on a reference interval, we continue traversing the path with LF-mapping, until we find the end of the interval. We add both sides of each node to the active set, again computing the appropriate distances.

##### 4.1.3 Extending the subgraph with snarls

If we want to extend the subgraph with snarls, we first need to find their boundary nodes. To do that, we iterate over all handles $x$ in the subgraph. If we are using a GBZ-base, we can simply use the `next` links in the node records. With a GBZ graph, we use a separate structure that has the links in a B-tree. If $next(x)$ is in the subgraph and edge $(x, next(x))$ would be in the canonical orientation, we add $(x, next(x))$ to the list of snarls. If $next(x)$ is outside the subgraph and we also want to include partially overlapping snarls, we similarly add $(x, next(x))$. If we want to include partially overlapping snarls but we did not find any snarls, we use breadth-first search to find a boundary node of a snarl containing the subgraph and add that snarl.

For each snarl $(x, next(x))$, we traverse the snarl using breadth-first search, starting from handles $x$ and $\overline{next(x)}$. We maintain active handles in a queue and visited handles in a hash set. When we visit an active handle $x$, we add the corresponding node to the subgraph in both orientations, if it is not already there. Then, for each edge $(x, y)$ to an unvisited handle $y$, we add both $y$ and $\overline{y}$ to the active handles.

###### 4.1.4 Extracting paths

Our path extraction algorithm is currently naive, as we are mostly interested in smaller subgraphs. First we decompress the path visits in the BWT fragments in the subgraph. For each handle $x$, we compute an array $A_x$ such that $A_x[i] = (y, j, \bot)$, where $LF(x, i) = (y, j)$. We store these arrays in a B-tree that maps $x$ to $A_x$. Then we iterate over the arrays. If we encounter $(y, j, \cdot)$ and handle $y$ is in the subgraph, we set $A_y[j] \leftarrow (\cdot, \cdot, \top)$ to indicate that the path visit $(y, j)$ has a predecessor in the subgraph. Then we do another pass over the arrays. If $A_x[i] = (y, j, \bot)$, we traverse the path starting from visit $(x, i)$ and collect it as a list of handles. If $z$ is the last handle and edge $(x, z)$ would be in the canonical orientation, we add the path to the list of paths in the subgraph.

The path deduplication algorithm is similarly naive. If the user wants to see only distinct local haplotypes, we sort the paths (lists of handles) in lexicographic order, remove duplicates, and store the number of duplicates in path metadata. Hash-based initial deduplication would be faster with large subgraphs, and we may implement it in the future.

###### 4.1.5 Computing the stable graph name

In order to compute a stable graph name (Supplement 3) for the subgraph, we need to list the nodes in order and the canonical edges adjacent to each node in the canonical order. This is the natural iteration order for both the subgraph data structure and GBZ. For each node identifier $v$ in sorted order, we first write a segment line with the identifier $v$ and label $\ell(v)$. Then we iterate over the edges in the record for the forward handle $\overrightarrow{x}$ and list the ones that are in the canonical orientation. Then we repeat the same for the reverse handle $\overleftarrow{x}$.

If the GBZ-base or the GBZ graph has a stable graph name stored in its metadata, we add a subgraph relationship between the subgraph and its parent graph. We also copy all known graph relationships from the parent graph.

###### 4.1.6 Aligning other paths to the reference

Some applications require CIGAR strings for the alignment (as encoded in the graph) between every other path and the reference path used in the query. This is currently done during serialization. GBZ-base version 0.7 will include an option for determining the alignments separately after a subgraph query.

In order to compute the alignment between two paths, we first find the longest common subsequence between the two lists of handles $A$ and $B$, with handle $x$ weighted by $|\ell(x)|$. We use Myers' $O(nd)$ algorithm [4] for this. Each matching handle $x$ becomes a match of length $|\ell(x)|$. For the diverging path fragments $A[i \dots j)$ and $B[i' \dots j')$ between two matching handles, we find the best-scoring alignment using only a mismatch, an insertion, and/or a deletion operation, considering only the lengths $|\ell(A[i \dots j))|$ and $|\ell(B[i' \dots j'))|$. While there could be base-level matches in the diverging parts, we assume that the graph construction pipeline had a reason for not aligning them.

#### 4.2 Querying a GAF-base

In order to extract alignments from a GAF-base, we need a source for node labels (sequences). A reference-free GAF-base can serve as a source. Otherwise we need a GBZ-base or a GBZ graph. If both the graph and the GAF-base store stable graph names, we use them for determining if the graph is a valid reference for the alignments. While we do not store the stable name of the subgraph in the read set, we include it if we serialize the alignments in GAF format.

When we continue a subgraph query with a GAF-base, we extract alignments overlapping with the subgraph into an in-memory read set data structure. The structure contains a list of alignment objects, which correspond to lines in a GAF file (Supplement 2). There is also a B-tree with a set of cached node records similar to the one in the subgraph data structure, as well as some metadata.

##### 4.2.1 Clustering the subgraph

As discussed in the main text, all alignments in a block $B$ with the handle interval $[a_B \dots b_B]$ overlapping that of the subgraph are candidates that may overlap with the subgraph. If there are large gaps between the node identifiers in the subgraph, the candidates may include many blocks with no alignments in the subgraph. To mitigate this issue, we cluster the subgraph into relatively dense intervals $[a_0 \dots b_0], [a_1 \dots b_1], \dots$. We currently start a new interval if the difference between successive node identifiers is more than 1000, but this may change in future releases.

##### 4.2.2 Finding candidates

For each interval $[a_i \dots b_i]$, we query table `Alignments` for blocks $B$ such that $[a_B \dots b_B] \cap [a_i \dots b_i] \neq \emptyset$. This uses a database index built for pairs $(a_B, b_B)$. The current query strategy appears to search for the last block $B$ such that $a_B \leq b_i$ and then traverse the index backwards, checking for overlaps. A two-dimensional index would traverse a smaller fraction of the index, but since we only expect around a million rows, this does not seem to be a bottleneck. To avoid duplicate work, we keep track of the database row identifiers we have already seen across all intervals.

##### 4.2.3 Decompressing a block

The actual decompression of a block into a list of alignments is relatively straightforward. After decompression, we have alignments, where the target path is represented by its GBWT starting position. To decompress the actual path, we iterate LF-mapping from the starting position until the end. We keep the alignment, if the path either overlaps with the subgraph or is fully contained in it, depending on query options.

We cache every node record we encounter while traversing the paths in the read set data structure. If the GAF-base is reference-free, we also get the sequence $\ell(x)$ for each handle $x$. Otherwise we first check if the subgraph also contains handle $x$ and copy the sequence from there. If not, we get the sequence from the parent graph.

If we chose to keep all alignments overlapping with the subgraph, we still have the option of clipping them to the subgraph (which is the default behavior). We output a separate alignment for each maximal fragment of the original alignment fully contained in the subgraph. To determine the query sequence interval corresponding to each fragment, we use the sequences $\ell(x)$ stored in the node records.

#### 5 PGGB graphs

While Minigraph–Cactus builds a pangenome graph by aligning other haplotypes to the selected reference, PanGenome Graph Builder (PGGB) aims to create unbiased graphs based on all-against-all alignment. Additionally, while the default Minigraph–Cactus graph has long unaligned segments clipped out, PGGB graphs are typically used as full graphs. This can cause issues when using PGGB graphs with tools such as GBZ-base designed for Minigraph–Cactus graphs.

The total length of the nodes in the Minigraph–Cactus graph we use is 3.75 Gbp, and the graph requires 5.7 GiB as a GBZ file (version 1) and 10.7 GiB as a GBZ-base. The corresponding numbers for the PGGB graph are 74.08 Gbp, 31.4 GiB, and 57.0 GiB. (Experimental version 3 of the GBZ format reduces the sizes to 3.35 GiB for the Minigraph–Cactus graph and 5.96 GiB for the PGGB graph, but it does not affect the size of the GBZ-base or the in-memory graph.) As the files are much larger, they are less convenient to use on a laptop.

There are also major differences in the structure of the graph. Minigraph–Cactus partitions the contigs by chromosome, creating a graph with 25 weakly connected components. PGGB does a true all-against-all alignment, resulting in a graph with 12,600 components. Most of the components appear to be unaligned contigs, while one massive component with 98% of the nodes represents all chromosomes aligned together. As there are no meaningful top-level chains, we cannot use top-level snarls in subgraph queries. Even if we chose to import nested snarls to GBZ-base, queries using them would be at a high risk of extracting a much larger subgraph than intended.

Beyond that, basic subgraph queries for a greedy context around a reference interval appear to work well with the PGGB graph. However, we have not tested them extensively.

Pangenome graph construction pipelines are typically developed together with tools using the graphs. Most of the properties of Minigraph–Cactus graphs result from attempts to make the graph a useful reference for mapping reads with Giraffe and calling variants and doing genome inference with various tools. There are currently no read aligners designed for PGGB graphs. While recent developments in distance indexing have made mapping reads with Giraffe to a PGGB graph computationally more feasible, the usefulness of such alignments in downstream applications remains an open question. As there is a lack of read alignments to PGGB graphs, we cannot currently say whether GAF-base can be used meaningfully with such alignments.

#### 6 GAF-base size breakdown

The major components of a GAF-base are the `Nodes` table storing the target paths and the binary blobs in table `Alignments`. Table 1 lists the size breakdown between those components for each

| Dataset | GBWT | Start | Names | Quality | Difference | Flags | Numbers | Optional |
|---|---|---|---|---|---|---|---|---|
| Element | 3.998 | 2.970 | 9.599 | 60.570 | 2.691 | 1.135 | 7.531 | 0.570 |
| Illumina | 3.965 | 2.184 | 5.737 | 7.372 | 2.484 | 0.773 | 3.688 | 0.442 |
| HiFi | 3.600 | 0.031 | 0.140 | 15.962 | 1.531 | 0.011 | 0.062 | 0.064 |
| ONT | 3.647 | 0.020 | 0.182 | 42.958 | 5.716 | 0.007 | 0.036 | 0.049 |

Table 1: GAF-base size breakdown, with the size of each component in gibibytes ($2^{30}$ bytes). Target paths stored in table `Nodes`, GBWT positions for path starts, read/pair names, quality strings, difference strings, binary flags, numbers, and optional fields.

dataset. For the target paths, we measure the size of the database file before and after inserting the data into the `Nodes` table. For the other components, we measure the total size of the binary blobs stored in the corresponding fields, excluding database overhead.

Quality strings are the largest single component for all datasets. For the other datasets except Illumina, they require more space than all the other components combined. The space used for difference strings depends on the error profile of the sequencing technology, being smallest with HiFi and largest with ONT. Target paths use similar space for all datasets, as the size of a GBWT index depends mostly on the number of node records. GBWT starting positions require 2 bytes/alignment in most cases. Read / pair names and numerical fields use substantial space with short reads and negligible space with long reads.

### 7 Subgraph queries

Full subgraph query results with a GBZ-base and a GBZ graph can be found in Figures 1 (with snarls) and 2 (without snarls). When we do not extend the subgraph with snarls, we avoid the outliers caused by large deletions within the greedy context. Apart from those outliers, query times are similar with and without snarls. Query performance is similar with Element and Illumina reads. Queries using ONT reads are somewhat slower than those with HiFi reads.

Queries for 100 bp intervals were much faster with a GBZ graph (mean 0.3 ms/query without reads) than with a GBZ-base (3 ms/query). Query times for such short intervals are dominated by the initial latency of retrieving the data. With 10 kbp intervals, the difference narrows down to 16 ms vs. 23 ms. Since retrieving more data from disk cache is almost as fast as using compressed in-memory data structures, in-memory computation dominates the overall query time.

As the subgraph query implementation was designed for GBZ-base, it creates an in-memory copy of the subgraph. With an in-memory GBZ graph, other approaches can be faster, depending on the intended uses for the subgraph. For example, vg uses subgraph overlays, which store a set of node identifiers and a pointer to the parent graph. The overlay implements a graph interface over the subgraph induced by the selected nodes by using the interface of the parent graph.

Peak memory usage for each query session can be found in Table 2. The memory usage of an individual query depends on the effective size of the subgraph (including the handles used by candidate alignments) and on the number and the length of the alignments in the subgraph. Peak memory usage is the maximum over all queries in the session.

Figure 3 shows the number of handles in the subgraph and in the candidate alignments with Illumina and HiFi reads. The latter is the effective size of the subgraph we must extract from GBZ-base

| Reads | Snarls | 100 bp | 1000 bp | 10000 bp | 100000 bp | 1000000 bp |
|---|---|---|---|---|---|---|
| None | None | 26.391 MiB | 37.594 MiB | 170.531 MiB | 1.832 GiB | 2.393 GiB |
| None | Contained | 26.266 MiB | 43.344 MiB | 321.422 MiB | 9.190 GiB | 2.282 GiB |
| Element | None | 56.594 MiB | 65.219 MiB | 207.969 MiB | 2.387 GiB | 3.200 GiB |
| Element | Contained | 57.500 MiB | 72.984 MiB | 370.500 MiB | 12.341 GiB | 3.152 GiB |
| Illumina | None | 58.172 MiB | 66.062 MiB | 200.391 MiB | 2.240 GiB | 2.900 GiB |
| Illumina | Contained | 59.328 MiB | 72.469 MiB | 359.438 MiB | 11.161 GiB | 2.932 GiB |
| HiFi | None | 178.297 MiB | 173.453 MiB | 310.594 MiB | 2.288 GiB | 2.652 GiB |
| HiFi | Contained | 178.062 MiB | 179.109 MiB | 452.703 MiB | 9.435 GiB | 2.692 GiB |
| ONT | None | 230.375 MiB | 242.031 MiB | 376.594 MiB | 2.040 GiB | 2.524 GiB |
| ONT | Contained | 229.062 MiB | 245.172 MiB | 509.406 MiB | 9.367 GiB | 2.498 GiB |

Table 2: Peak memory usage (peak resident set size) for each query session when using a GBZ-base. Memory usage with a GBZ graph is approximately 12 GiB higher.

when we want to extract the alignments overlapping with the subgraph. With 100 bp queries, this effective subgraph is orders of magnitude larger than the actual subgraph. This explains why there is little difference in query times between 100 bp and 10 kbp queries when we also extract alignments from a GAF-base.

### 8 Specific commands used

We assume that the following directories are located on a fast local SSD:

- `$WORK`: Working directory containing the graphs and the reads
- `$TMPDIR`: Temporary directory
- `./logs`, `./db-logs`, and `./gbz-logs`: Logs

All relevant tools are assumed to be found in the path. GBZ-base / GAF-base tools are compiled with:

```
cargo build --release --features benchmark
```

Variable `$1` in the scripts is the name of the dataset (e.g. `illumina`).

#### 8.1 GBZ-base construction

We copied the default graph `default.gbz` to the current directory. Then we build GBZ-base with:

```
LOGFILE=logs/gbz-base-${1}.log
rm -f $LOGFILE

gbz2db --overwrite ${1}.gbz 2>> $LOGFILE
```

The log contains wall-clock time and peak memory usage for the construction, as well as the size of the GBZ-base database.

#### 8.2 GAF-base construction

##### 8.2.1 Mapping reads with Giraffe

We use the following script for mapping the reads:

```
if [ "$1" == element ]; then
  GRAPH=${WORK}/default.gbz
  KMERS=${WORK}/${1}.kff
  SAMPLING="--haplotype-name ${WORK}/default.hapl --kff-name $KMERS"
  PAIRED="-i"
elif [ "$1" == illumina ]; then
  GRAPH=${WORK}/default.gbz
  KMERS=${WORK}/${1}.kff
  SAMPLING="--haplotype-name ${WORK}/default.hapl --kff-name $KMERS"
  PAIRED="-i"
elif [ "$1" == hifi ]; then
  GRAPH=${WORK}/filtered.gbz
  PRESET="--parameter-preset hifi"
elif [ "$1" == ont ]; then
  GRAPH=${WORK}/filtered.gbz
  PRESET="--parameter-preset r10"
fi

LOGFILE=logs/giraffe-${1}.log
rm -f $LOGFILE

INPUT=${WORK}/${1}.fq.gz
OUTPUT=${WORK}/${1}.gaf.gz
THREADS=32

if [ "$KMERS" != "" ]; then
  kmc -k29 -m128 -okff -t${THREADS} -hp $INPUT ${WORK}/${1} $TMPDIR
fi

vg giraffe -p -t $THREADS -Z $GRAPH $SAMPLING $PAIRED -f $INPUT $PRESET -o gaf \
  2>> $LOGFILE | bgzip > $OUTPUT
```

With short reads (Element and Illumina), we do paired-end mapping to a personalized graph. We count $k$-mers separately with `kmc` and let `vg giraffe` run haplotype sampling and build indexes for the personalized graph. While `vg giraffe` could also run `kmc` automatically, there is a bug in vg version 1.75.0 that makes the alignment output incorrect when option `-p` / `--progress` is used. This bug has been fixed in vg version 1.75.1.

With long reads (HiFi and ONT), we use the appropriate preset and map the reads to the frequency-filtered graph. `vg giraffe` builds the indexes automatically if they do not exist. This assumes that the datasets are mapped one at a time. If both datasets are mapped concurrently, the indexes should be built first with `vg autoindex` to avoid conflicts between multiple jobs building the same indexes.

##### 8.2.2 GAF sorting

We use the following script for sorting the original GAF file:

```
if [ "$1" == element ]; then
  SORT_BLOCK=1M
  DB_BLOCK=1000
elif [ "$1" == illumina ]; then
  SORT_BLOCK=1M
  DB_BLOCK=1000
elif [ "$1" == hifi ]; then
  SORT_BLOCK=10k
  DB_BLOCK=10
elif [ "$1" == ont ]; then
  SORT_BLOCK=10k
  DB_BLOCK=10
fi

LOGFILE=logs/gaf-base-${1}.log
rm -f $LOGFILE

INPUT=${WORK}/${1}.gaf.gz
OUTPUT=${WORK}/${1}.sorted.gaf.gz
DB=${WORK}/${1}.db
GRAPH=${WORK}/default.gbz
DECOMPRESSED=${WORK}/temp.gaf.gz
THREADS=16
BGZIP_THREADS=6

gafsort -p -r $SORT_BLOCK -t $THREADS $INPUT 2>> $LOGFILE \
  | bgzip --threads $BGZIP_THREADS > $OUTPUT
```

The number of sorting threads primarily affects the intermediate merging rounds. The final merge is mostly sequential, while a few threads are already enough to saturate the gzip decompression in `gafsort` during the initial sorting. Wall-clock time and peak memory usage can be found in the log.

##### 8.2.3 GAF-base construction

The script continues with GAF-base construction:

```
echo >> $LOGFILE

gaf2db --overwrite -b $DB_BLOCK -o $DB $OUTPUT 2>> $LOGFILE
```

Wall-clock time and peak memory usage can be found in the log again. The construction also reports the size of the database after creating each table, as well as the total sizes of various fields in table `Alignments`. GAF-base size breakdown (Supplement 6) can be derived from this information.

##### 8.2.4 GAF-base decompression

The final part of the script is GAF-base decompression:

```
echo >> $LOGFILE

(db2gaf -r $GRAPH $DB 2>> $LOGFILE) \
  | bgzip --threads $BGZIP_THREADS > $DECOMPRESSED
rm -f $DECOMPRESSED
```

Wall-clock time and peak memory usage can be found in the log. We need a reference graph for converting the internal representation of the difference strings to actual difference strings that in some cases contain the reference bases corresponding to the edit operations.

#### 8.3 GAF-base size comparison

We determine the file sizes for each dataset using the following script:

```
if [ "$1" == element ]; then
  READ_LENGTH="--read-length short"
elif [ "$1" == illumina ]; then
  READ_LENGTH="--read-length short"
elif [ "$1" == hifi ]; then
  READ_LENGTH="--read-length long"
elif [ "$1" == ont ]; then
  READ_LENGTH="--read-length long"
fi

LOGFILE=logs/file-sizes-${1}.log
rm -f $LOGFILE

BASENAME=${WORK}/${1}
GRAPH=${WORK}/default.gbz
REFERENCE=${WORK}/chm13.fa
REFERENCE_SAMPLE=CHM13
THREADS=32

# Extract the correct reference, if necessary.
if [ ! -f $REFERENCE ]; then
  vg paths -x $GRAPH -S $REFERENCE_SAMPLE -F > $REFERENCE
  samtools faidx $REFERENCE
fi

# Compressed GAF
ls -l ${BASENAME}.gaf.gz >> $LOGFILE

# Sorted compressed GAF
ls -l ${BASENAME}.sorted.gaf.gz >> $LOGFILE
```

```
# GAF-base
ls -l ${BASENAME}.db >> $LOGFILE

# GAF
bgzip -d -c --threads $THREADS ${BASENAME}.gaf.gz > ${BASENAME}.gaf
ls -l ${BASENAME}.gaf >> $LOGFILE
rm -f ${BASENAME}.gaf

# GAM
vg convert -t $THREADS --gaf-to-gam ${BASENAME}.gaf.gz $GRAPH > ${BASENAME}.gam
ls -l ${BASENAME}.gam >> $LOGFILE
rm -f ${BASENAME}.gam

# BAM
vg surject -t $THREADS -x $GRAPH --into-ref $REFERENCE_SAMPLE $READ_LENGTH \
  --gaf-input --bam-output ${BASENAME}.gaf.gz > ${BASENAME}.bam
ls -l ${BASENAME}.bam >> $LOGFILE

# Sorted CRAM (from BAM)
samtools sort --threads $THREADS -m 4G -T ${TMPDIR}/samtools ${BASENAME}.bam \
  > ${BASENAME}.sorted.bam
samtools view --threads $THREADS --reference $REFERENCE \
  --cram ${BASENAME}.sorted.bam > ${BASENAME}.cram
ls -l ${BASENAME}.cram >> $LOGFILE
rm -f ${BASENAME}.bam ${BASENAME}.sorted.bam ${BASENAME}.cram
```

We convert GAF to GAM with `vg convert` and GAF to BAM with `vg surject`. Then we obtain sorted CRAM from BAM by sorting it with `samtools sort` and converting the result with `samtools view`.

Since we used SAMtools 1.19.2 with default parameters, the CRAM files were of version 3.0. CRAM 3.1 files would have been smaller due to the availability of better codecs.

##### 8.3.1 Subgraph queries

In order to do snarl-based queries with an in-memory GBZ graph, we had to extract top-level chains (`default.chains`) from a distance index:

```
# Build a minimal distance index
vg index --no-nested-distance -j default.dist default.gbz

# Extract top-level chains
vg chains -p -o default.chains default.gbz default.dist
```

For benchmarking subgraph queries, we copied all GAF-bases to the current directory. Before each session, we purged disk caches with `sudo purge`. We ran `query-benchmark` with the following options:

- `--gbz-base default.gbz.db` (when using GBZ-base) or `--gbz default.gbz` and `--chains` `default.chains` (when using GBZ)

- With no GAF-base and with `--gaf-base {$1}.db` for each dataset `$1`
- Using CHM13 as the reference: `--faidx CHM13.fa.fai --sample CHM13`
- `--interval-length $N` for `$N` in $\{100, 1000, 10000, 100000, 1000000\}$
- `--num-queries $M`, with `$M` as 1000 otherwise and as 100 when `$N` was 1000000
- With and without `--snarls`
- `--verbose`

We redirected the standard output to a log file. For each query, the log contains a line with the following TAB-separated fields:

1. Description of the query.
2. Number of nodes in the subgraph.
3. Number of alignment fragments in the output.
4. Number of original alignments in the subgraph.
5. Number of alignment blocks decompressed.
6. Number of candidate alignments in the blocks.
7. Number of distinct handles in the candidates.
8. Query time in seconds.

#### 9 AI disclosure statement

Initial versions of various functionalities in the code and data processing scripts were implemented by Claude Code, using various versions of the Opus and Sonnet models. Manual code editing was assisted by code completions from GitHub Copilot. No AI tools were used for generating or editing the text and figures in the manuscript and its supplementary materials.

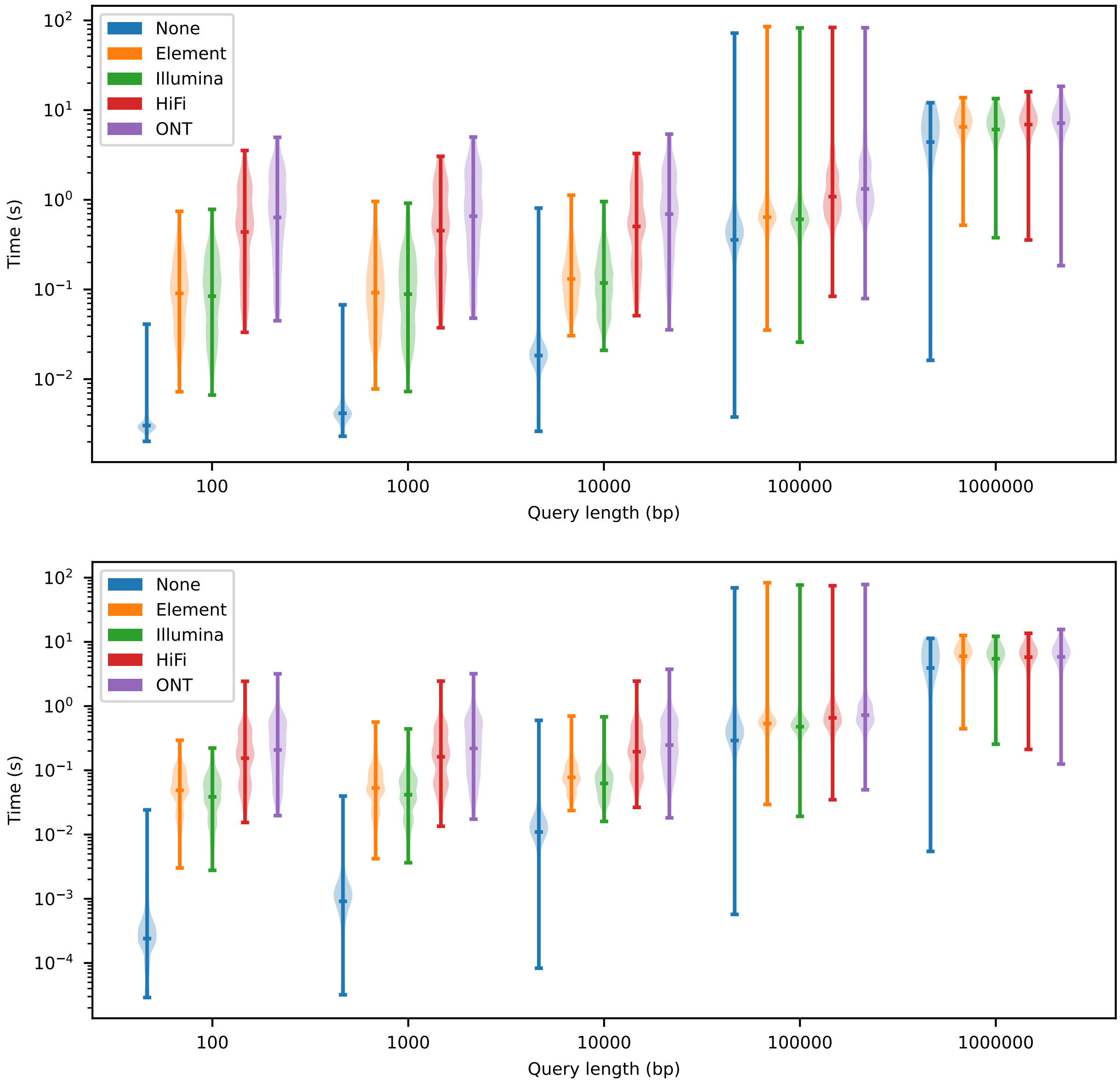


Figure 1: Violin plots for subgraph query times with random CHM13 intervals, 100 bp greedy context, and contained snarls. Queries with a GBZ-base (top) and with an in-memory GBZ graph (bottom). Subgraphs with no reads and with various GAF-bases.

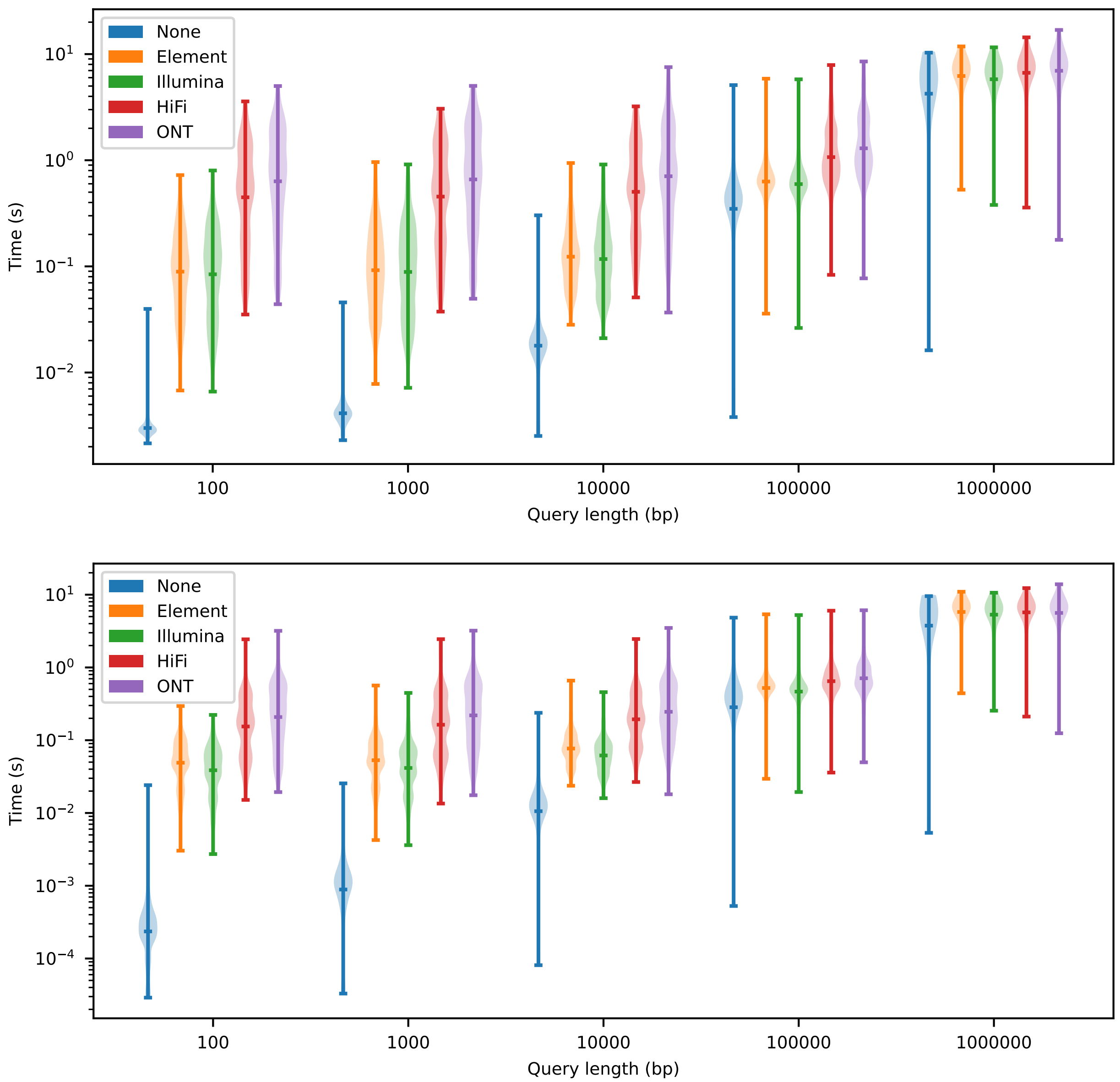


Figure 2: Violin plots for subgraph query times with random CHM13 intervals, 100 bp greedy context, and no snarls. Queries with a GBZ-base (top) and with an in-memory GBZ graph (bottom). Subgraphs with no reads and with various GAF-bases.

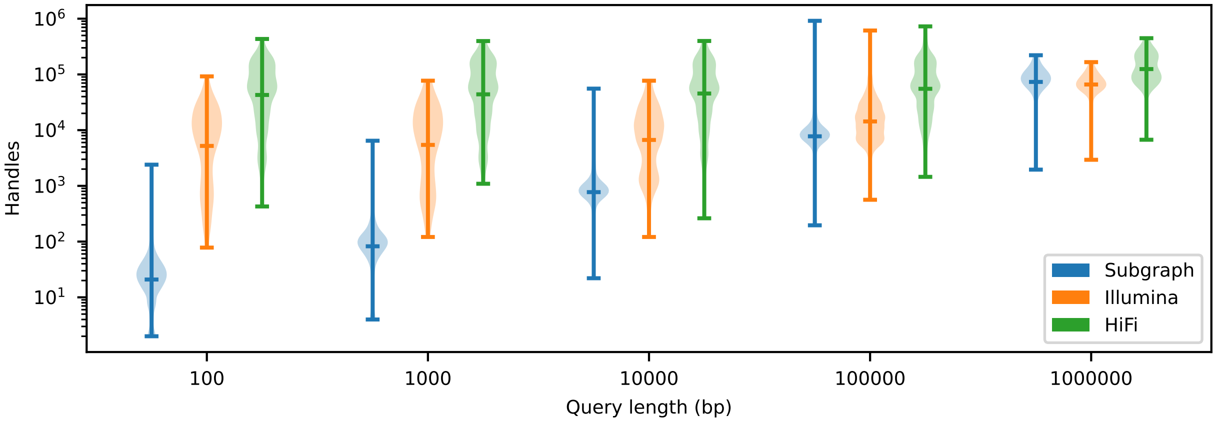


Figure 3: Violin plots for the number of handles in the subgraph and in the candidate alignments of Illumina / HiFi reads with random CHM13 intervals, 100 bp greedy context, and contained snarls.
